# Automated quantitative trait locus analysis (AutoQTL)

**DOI:** 10.1101/2023.01.12.523835

**Authors:** Philip J. Freda, Attri Ghosh, Elizabeth Zhang, Tianhao Luo, Apurva Chitre, Oksana Polesskaya, Celine L. St. Pierre, Jianjun Gao, Connor D. Martin, Hao Chen, Angel G. Garcia-Martinez, Tengfei Wang, Wenyan Han, Keita Ishiwari, Paul Meyer, Alexander Lamparelli, Christopher P. King, Abraham A. Palmer, Ruowang Li, Jason H. Moore

**Author notes:** Equal Authorship (Philip J. Freda, Attri Ghosh). Corresponding Author –.

## Abstract

**Background:** Quantitative Trait Locus (QTL) analysis and Genome-Wide Association Studies (GWAS) have the power to identify variants that capture significant levels of phenotypic variance in complex traits. However, effort and time are required to select the best methods and optimize parameters and pre-processing steps. Although machine learning approaches have been shown to greatly assist in optimization and data processing, applying them to QTL analysis and GWAS is challenging due to the complexity of large, heterogenous datasets. Here, we describe proof-of-concept for an automated machine learning approach, AutoQTL, with the ability to automate many complex decisions related to analysis of complex traits and generate diverse solutions to describe relationships that exist in genetic data.

**Results:** Using a dataset of 18 putative QTL from a large-scale GWAS of body mass index in the laboratory rat, *Rattus norvegicus*, AutoQTL captures the phenotypic variance explained under a standard additive model while also providing evidence of non-additive effects including deviations from additivity and 2-way epistatic interactions from simulated data via multiple optimal solutions. Additionally, feature importance metrics provide different insights into the inheritance models and predictive power of multiple GWAS-derived putative QTL.

**Conclusions:** This proof-of-concept illustrates that automated machine learning techniques can be applied to genetic data and has the potential to detect both additive and non-additive effects via various optimal solutions and feature importance metrics. In the future, we aim to expand AutoQTL to accommodate omics-level datasets with intelligent feature selection strategies.

## Background

Quantitative trait locus (QTL) analysis and genome-wide association studies (GWAS) have collectively revolutionized the analysis of complex traits, including human disease. However, these methods commonly rely on univariate or interval mapping approaches and an additive genetic model to capture phenotypic variance [1]. Thus, they are limited in their ability to model non-additive genotypic effects, including epistatic interactions [2]. However, there are significant exceptions to this general trend.

A recent large-scale study in *Saccharomyces cerevisiae* identified that of the 61% broad-sense heritability detected, 21% was due to non-additive effects (dominance deviations and epistasis) [3]. In a related previous study, also in *S. cerevisiae*, of the 91.7% of the phenotypic variance attributed to genetic sources, non-additive effects accounted for 18.7% [4] Furthermore, non-additive QTL outnumbered additive QTL 3:1. These studies illustrate that significant sources of non-additive genetic variation, which are not usually detected using standard approaches, exist in living systems. However, in both studies, the accounting for non-additive genetic effects was achieved by constructing a rigorous breeding design coupled with thousands of replicated lineages. This level of structure in known relatedness and replication can be especially challenging in human genetic studies or in non-model systems.

An alternative, and potentially more exploratory, approach is to employ automated machine learning (autoML) techniques, which can be designed to detect both additive and non-additive effects while reducing the need to invest significant resources in constructing specific methodologies and selecting specialized parameters. AutoML approaches have the advantage of exploring vast parameter spaces while performing both algorithm selection and hyperparameter optimization without express human interaction post-initialization and can therefore be unleashed to detect patterns in large and complex datasets. Even though, to date, autoML packages specifically designed for phenotype to genotype association in omics level data have not been fully explored, there have been several significant and promising attempts. As initial examples, there have been studies that have identified epistatic interactions using random forest (RF) methodologies for major adverse cardiovascular events [5] and Alzheimer’s Disease [6] from simulated and real-world GWAS-scale data. RF-based methodologies have also been designed to identify non-coding variants [7,8] and to predict rare variant pathogenicity [9]. Examples proximal to GWAS-level exploratory analyses come from successful applications of the software package TPOT [10–12] to several omics datasets built to explore medically relevant phenotypes. TPOT, the Tree-based Pipeline Optimization Tool, selects Pareto optimal pipelines with the highest multi-objective fitness after a user designated number of genetic programming (GP) generations. In this way, TPOT can optimize algorithms and hyperparameters using an approach akin to Darwinian evolution. To date, TPOT has been applied to RNAseq data to identify pathways and genes associated with major depressive disorder [12] and schizophrenia [13] as well as to metabolomics data to study type 2 diabetes [14] and coronary artery disease [15]. As for genotype to phenotype association, TPOT was recently used to analyze a large cohort from the UK Biobank to study coronary artery disease using an expert-knowledge feature filter [16]. A subset of 28 SNPs was identified and were linked to putative genes related to atherosclerotic plaques and myocardial infarction.

Despite the promising examples explored above, the application of autoML approaches to QTL analysis and GWAS has historically been limited for two reasons. The first is best described as the *curse of dimensionality* where the number of predictors (features or variants) are much larger than the number of observations. To remain computational tractable, autoML approaches to omics-level data must parse and filter predictors based on expert biological knowledge or computationally derived methods to reduce dimensionality. However, optimizing these strategies can prove to be difficult, especially when it comes to universality of the approach. Second, the vast majority of autoML software packages are not specifically designed with quantitative genetics and epistasis as a core part of algorithm and hyperparameter optimization. Designing an approach specifically centered around the genetic analysis of complex traits with these considerations in mind has the capacity to elucidate aspects of biology not frequently explored by standard genetic analyses.

Here, we provide a proof-of-concept for an open-source automated machine learning (AutoML) method and software, AutoQTL, to automate QTL analysis by building an analytics pipeline optimized for explaining variation in a complex trait given a set of genetic variants. Most AutoML methods are agnostic to the application domain and thus unaware of decisions important to geneticists such as selecting features by allele frequency or feature encoding to capture recessive or dominant effects. Thus, we have developed AutoQTL as a method to automatically explore important aspects of quantitative genetics and the analysis of complex traits. Central to this method is an algorithm designed to make decisions a geneticist might make when planning and executing QTL analysis. These decisions include genetics-based encoding of the genotypes for each genetic variant (i.e., feature encoding), selection of genetic variants based on genetic principles such as allele or genotype frequency (i.e., feature selection), and the selection of a parametric statistical or machine learning based regression method for relating genotype to phenotype, allowing AutoQTL to explore all sources of variation, including non-additive effects.

## Methods

Key to AutoQTL is the data structure for representing QTL analysis pipelines for optimization. We utilize expression trees as an intuitive graph-based data structure for representing pipelines where a regression method is selected in the root node, feature encoding and feature selection methods can be selected for the child nodes, and hyperparameters selected as terminals of the tree.

We selected GP as the search and optimization method for AutoQTL given it is designed to generate, diversify, evaluate, and select optimal expression trees given some fitness or quality metrics [17–19]. Below, we describe the components of AutoQTL pipelines, GP optimization, the data, and the experiments used to evaluate the method.

### AutoQTL Operator Classes

All operators (root nodes and child nodes) have been implemented using the Python programming language [20] including existing implementations in the scikit-learn [21] machine learning package. The three operator classes for AutoQTL are feature encoders, feature selectors, and regression methods.

We have implemented operators for the encoding phase that demonstrate an additive inheritance model and alternatives [22] (e.g. dominance deviations and heterosis). These operators modify the original values of the genotypes (Figure 1A) into forms that can express varied genetic relationships, allowing our program to introduce inheritance models beyond those used in standard QTL analyses. Currently, there are five different encoding operators which are applied across all genotypes in the data outside of a standard additive model (default encoding where AA = 0, Aa = 1, and aa = 2). Figure 1B illustrates examples of encoding changes. For a complete list of all the five additional encoding operators, refer to File S1. If a pipeline contains two or more encoding operators, multiple transformations will occur in a sequence to achieve inheritance models not necessarily indicative of the initial encoder and can even lead to final encodings not possible via one encoder (examples in File S1).

**Figure 1.**
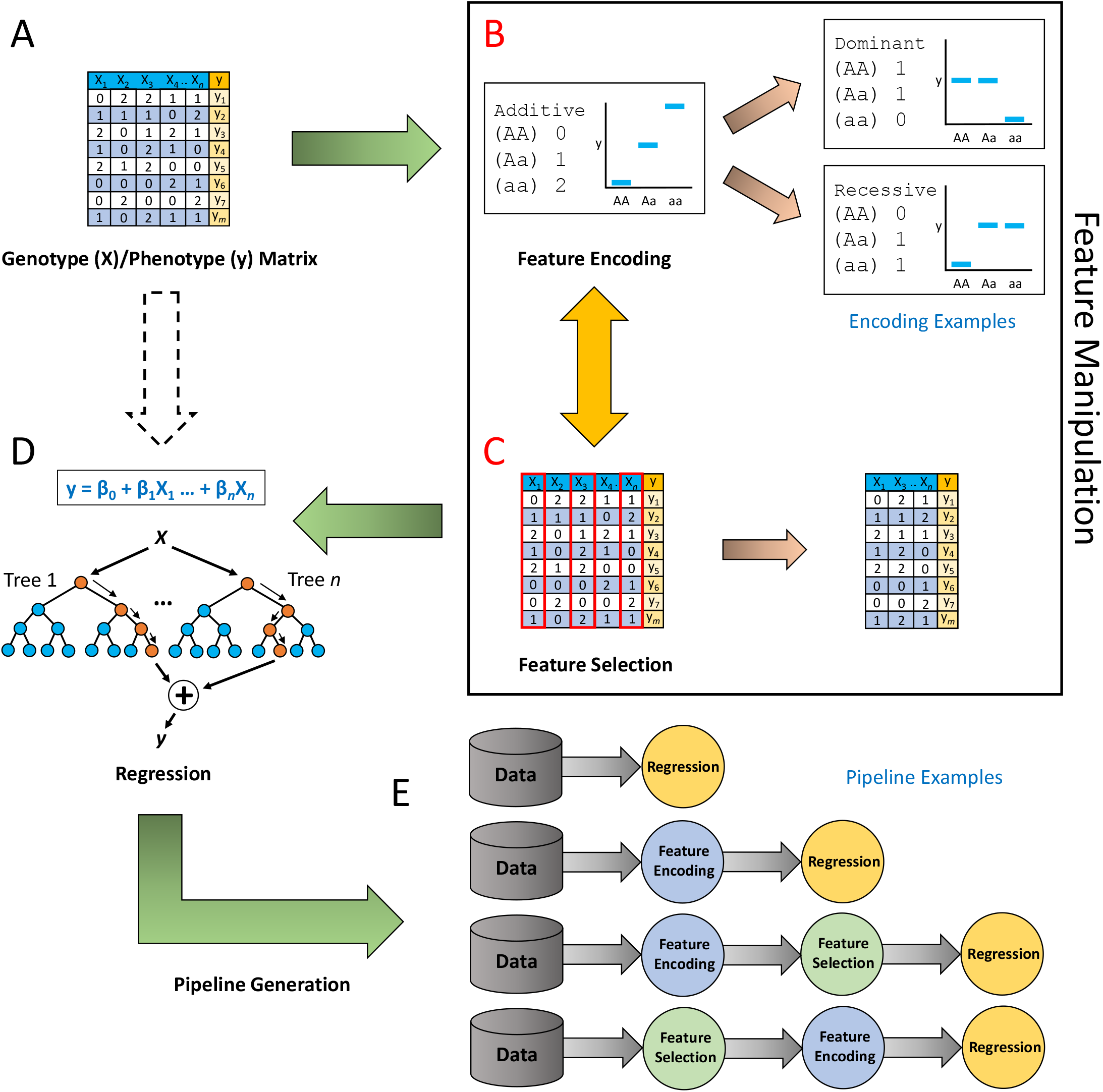
Conceptual image of AutoQT’s workflow. A. A genotype/phenotype matrix is read into AutoQTL. B. An optional feature encoding step recodes the data into five possible and distinct genetic models (File S1). C. An optional feature selection step where features (loci) are removed by a selection operator and hyperparameter. Note that feature encoding and feature selection steps are optional and can occur more than once in any order. D. A root regressor and hyperparameters (machine learning) are selected. E. Examples of pipelines that are scored through GP.

In the feature selection phase, we remove features based on pre-designed criteria. AutoQTL has three feature selection operators with their own respective hyperparameter ranges and increments: (Figure 1C.) 1.) The scikit-learn VarianceThreshold (VT) operator removes features that do not satisfy a minimum variance threshold (0 – 0.35, increments of 0.05), 2.) the scikit-learn SelectPercentile (SP) operator selects the top *n* percentile of features based on an F-regression R^2^ score (where *n* can be between 5% – 95% in increments of 5%), and 3.) the genotype frequency (GF) operator removes features if any genotype frequency is below a threshold (0 – 0.35, increments of 0.05). Like with feature encoding, multiple feature selection phases may be present in an AutoQTL pipeline, leading to combinatorial effects of feature selection based on multiple criteria and hyperparameters.

We have implemented three regression method operators from the scikit-learn package which serve as the root node of a pipeline: LinearRegression (LR), and two machine learning regressors: DecisionTreeRegressor (DT) (individual tree-based model) and RandomForestRegressor (RF) (ensemble tree-based model). Unlike feature encoding and feature selection phases, only one regression (root) can be selected per pipeline. The phenotype explored in this proof of concept is continuous, but we will expand AutoQTL to also handle discrete phenotypes by adding classifier methods readily available in scikit-learn.

### Pipeline Generation via Genetic Programming

Graph-based data structures are nonlinear data structures consisting of two primary components: nodes and edges [23]. Each edge connects two nodes to form a path that represents a relationship between two entities. Expression trees are a type of graph-based data structure where the root node along with each internal node refers to an operator and each terminal node is an operand [23] (see File S1 for example). In this work, we represent our machine learning pipelines as expression trees. This form of representation allows diverse solutions due to the flexibility of a tree structure. Nodes can be easily added, deleted, or modified in the expression tree allowing our program to explore various possible solutions and evolve to generate the best possible machine learning pipelines for QTL analysis. GP is a common approach for manipulating and optimizing expression trees [17–19]. Adapting concepts from modern genetics and Darwinian natural selection, GP has the capability to improve pipelines after each successive generation of selection. Hence, we selected GP as our search and optimization technique due to its unique design for expression trees and ability to select optimal pipelines.

We have adapted the core algorithm comprised of GP and Pareto optimization [24] from the python package TPOT (Tree-based Pipeline Optimization Tool) [10,12,25]. AutoQTL pipelines are generated using GP implemented in DEAP (Distributed Evolutionary Algorithms in Python) [26].

We split the input data randomly into 80%/20% split and use the 20% split data as a holdout dataset to evaluate our final Pareto front pipelines at the end of the process to assess for overfitting. We then split the remaining input data in half to produce the training and testing sets - each containing 40% of the original data. AutoQTL pipelines are generated randomly in the first generation and are each assigned a fitness score. Each AutoQTL pipeline is evaluated using two metrics: test split R^2^ and the difference score (DS), which is calculated as: 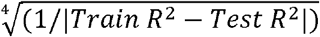. The DS is transformed using a biquadratic root to reduce the scale. The objective of DS is to penalize overfitting. Thus, test R^2^ and DS are the objectives of Pareto optimization performed at the end of each generation. Maximizing both metrics allows for more varied Pareto fronts with multiple optimal solutions. Once all the pipelines are evaluated, the Pareto front is updated.

To create the next generation, new pipelines (offspring) are generated from the existing population of pipelines by applying mutation and crossover. Those pipelines with higher fitness contribute more phases (akin to genes in biological evolution) of their pipelines compared to pipelines of lower fitness. Mutation of a pipeline can be performed in the following three ways, and each has equal probability of occurring: 1.) Uniform mutation: Either a random AutoQTL operator in a pipeline is replaced with a new randomly selected operator (e.g., an additive encoder switching to a dominance encoder) or the hyperparameter values of an operator is modified (e.g., variance threshold of 0.05 changed to 0.10). 2.) Insert mutation: A new randomly selected AutoQTL operator is inserted in a pipeline at a random position, adding a step to a pipeline. 3.) Shrink mutation: A random AutoQTL operator is removed from the pipeline. Examples of mutation events are found in File S1.

Crossover is performed by splitting two eligible chosen pipelines at a random point in the pipeline tree and then swapping the subtrees. Eligibility between two pipelines is defined when both pipelines have a phase operator in common (e.g., both have a feature encoding phase). This is known as one-point crossover. This produces two offspring pipelines. The first offspring is selected to populate the next generation while the second is discarded. An example of a crossover event is found in File S1.

Once offspring are created, a new population for the next generation is produced by selecting pipelines from the present parent population and offspring population with the highest fitness by the selection algorithm Non-dominated Sorting Genetic Algorithm (NSGA)-II [27]. This selection process evolves the population by retaining pipelines with optimal fitness. Each new population is evaluated, and the Pareto front is then updated. This sequence of pipeline evolution continues for *n* number of generations set by the user. At the end of *n^th^* generation, the Pareto front contains the most optimal pipelines discovered by AutoQTL over *n* generations.

For our analyses, population size was set at 100, offspring size was set at 100, the mutation rate was set at 90%, and crossover rate was set at 10% for a maximum of 25 generations. All these parameters can be changed by the user before initializing AutoQTL.

### Feature Importance

To determine each feature’s (locus) importance to each pipeline, we utilized Shapley [28] values. Shapley values are local metrics derived from game theory that correspond to how much each feature contributes to a ML model prediction. The SHAP (SHapley Additive exPlanations) package in python was used to calculate Shapley values for each feature in Pareto optimal pipelines at the end of the GP process. In a case where a feature selector was selected as a node in a Pareto optimal pipeline and a feature was removed, its Shapley value is 0 as this feature is not contributing to that pipeline’s model and, thus, has no predictive value. Because GWAS and QTL analyses rely on beta coefficients and *p*-values to determine variant effect size and significance, we incorporated a feature importance procedure to compare and rank features by measuring their model contributions. Although not analogous to standard hypothesis-driven statistical metrics used in GWAS and QTL analysis, Shapley values allow the user to select putative QTL (highly predictive features) for further study.

### Benchmark Data

To test AutoQT’s GP pipeline processing and optimization, we used genotype data for 18 putative QTL (significance threshold = –log10 *p* > 5.6) for body mass index, defined as the distance from the tip of the nose to the tip of the tail, in rats (*Rattus norvegicus*) from a genome-wide association study (GWAS) using 3,400,759 single nucleotide polymorphisms (SNPs) [29,30]. We used these 18 putative QTL to test the capability of AutoQTL to first capture phenotypic variance in a subset of loci before expanding to a larger, exploratory analysis. Additionally, a smaller dataset allows us to determine if AutoQTL can detect interactions among random noise or features with main effects. Quantile normalized residuals for 5,566 rats were corrected for relatedness between individuals by performing a principal components analysis (PCA) in PLINK [31] using R [32]. A multiple linear regression between the residuals and the first 10 principal components was performed in R. Residuals were then retained as the phenotype values for all further analyses.

### Validation of AutoQTL

The multiple LR R^2^ (0.0968) for the genotype/phenotype dataset from the 5,566 rat cohort for the 18 putative QTL [29,30] was determined in Python using LR class from scikit-learn package outside of AutoQTL to determine the total phenotypic variance (V_*P*_) explained by the linear model. Since the 18 putative QTL were identified using standard GWAS approaches (linear regression assuming an additive inheritance model), AutoQTL should generate a final Pareto optimal pipeline with linear regression (LR) as the only phase in each run that captures all the V_*P*_ (R^2^) in the dataset. However, since AutoQTL randomly splits the data, we compared each pipeline’s test R^2^ to the R^2^ of the test split with basic LR since it will vary from the LR R^2^ of the full dataset (unsplit). We performed 10 replicates, each with a distinct random seed, ensuring different data splits, and determined if one of the Pareto optimal solutions included LR as the only pipeline phase and captured the entirety of the V_*P*_ in the test set for that split. We also explored the other Pareto optimal solutions and compared them to LR alone.

### Detecting Epistasis with AutoQTL

To evaluate AutoQT’s ability to detect genetic epistasis, an interaction between two or more genes in which the effect of one gene is modified (e.g. masked, inhibited, or suppressed) by one or more genes [33], we created 2-way interactions using the XOR penetrance function (File S1), which leads to a marginal penetrance of 0.5 assuming equal allele frequencies under Hardy-Weinberg equilibrium for every two-locus genotype combination [34,35]. We generated a total of nine two-way interactions with each of the 18 loci involved in a two-way interaction with one other loci without overlap (e.g., each 2-way interaction is unique, and no loci were used more than once). To achieve this, we matched two-locus genotype combinations that result in a penetrance of 1 in the XOR model to the top 50% (in ascending order) of phenotypic scores (y). This results in the strongest possible XOR signal. However, to generate a total R^2^ matching that of the 18 QTL dataset (0.0968) detectable by AutoQTL’s machine learning regressors, we randomly shuffled a percentage of the genotypes to weaken the total signal, partially breaking the strong association of complete penetrance with the phenotype (see File S1 for explanation and File S2 for R code). We did this for each 2-way interaction until we achieved a total V_*P*_ (R^2^) detectable in AutoQTL by machine learning regressors of approximately 0.10 with all nine interactions. As a result of this shuffling, all univariate effects totaled nearly zero V_*P*_ (multiple LR R^2^ after shuffling = 0.00480). Thus, the distribution of genotype and allele frequencies remains intact while we alter the genotype-phenotype relationships of the dataset by manipulating two loci at a time (File S2). We inputted this dataset into AutoQTL and compared Pareto optimal pipelines to those of the 18 putative QTL experiment. Our hypothesis is that, in the presence of interactions, AutoQTL will select a higher proportion of machine learning regression roots (e.g., RF and DT) versus LR in the final Pareto front. This is because RF and DT have been shown to detect interactions whereas multiple LR models without interaction terms cannot [36,37]. Furthermore, cartesian products (multiplicative interaction terms) cannot accurately describe penetrance functions as the interaction effect is not a product of the genotypic states but rather described in marginal penetrances (in this case, pure 2-way epistasis in XOR, either 1 or 0 [35]). We performed 10 replicates, with the same random seeds as the 18 QTL validation experiment.

### Describing Phenotypic Variance as Interactions Increase

To determine how V_*P*_ explained by LR and machine learning regressors changes in the presence of an increasing quantity of 2-way epistatic interactions, we reduced the significant main effects from the original 18 putative QTL dataset by shuffling columns (File S1; File S2) as we did in the previous XOR experiment. Then we added 2-way XOR interactions, one at a time, and entered each dataset (ranging from 0-9 interactions) into AutoQTL. As with in the XOR dataset experiment, each of the 18 loci were involved in a two-way interaction with one other loci without overlap. This was performed for 10 replicate datasets per interaction. This resulted in 100 datasets, 10 with no interactions, and 90 with interactions ranging from one to nine. For this dataset, we generated interactions with a maximum test R^2^ detectable by machine learning regressors of approximately 0.15 with all nine interactions to better illustrate the difference between regressor types.

To illustrate how V*_P_* explained changes with increasing interactions in the presence of main effects, we also replicated the above procedure by creating 2-way interactions, one at a time, while keeping the original main effects of all other loci intact. Each time a new interaction pair was generated, two loci were selected and had their main effects converted into a 2-way interaction, thereby breaking the original main effect. In other words, as the number of interactions increased, the number of loci with significant main effects reduced. To accurately compare between test the LR R^2^ with 18 main effects (18 QTL validation experiment) and with nine interactions, the maximum test R^2^ explained by 9 interactions via machine learning regressors was limited to 0.10. We performed this experiment with ten replication datasets per interaction for a total of 100 datasets. We also generated Shapely feature importance for an example run (R random seed = and AutoQTL random seed = 0) across interactions from zero to nine to explore how the difference in feature importance between interacting loci is altered as main effects are replaced by interactions.

## Results

### AutoQTL captures Phenotypic Variance of GWAS QTL and Detects Dominance Deviations

All the AutoQTL final Pareto fronts for each separate random seed generated a pipeline (pipeline marked with a star in Figure 2A) where only the root LR method was selected as the only operator resulting in the test R^2^ matching that of the test R^2^ before GP was executed in AutoQTL (File S3). This validates AutoQT’s ability to capture the V_*P*_ explained by multiple LR alone. However, in addition to this pipeline, each final Pareto front also includes pipelines with lower R^2^ than LR alone but with feature selection and/or encoding operators selected resulting in a higher DS (smaller difference in R^2^ between training and testing). Table 1 highlights the pipelines in an example run of AutoQTL (random seed = 12) that are also illustrated in Figure 2A. The two pipelines that are optimized in both metrics (Pareto efficient), which are in the middle of Figure 2A with arrows pointing to them, are in bold. All Pareto optimal pipelines over the six generations for this example run are found in File S4. On average, final Pareto fronts for the 18 QTL dataset over the ten runs contain 7 pipelines and 88.5% of these pipelines had LR selected as the root regressor (File S5).

**Figure 2.**
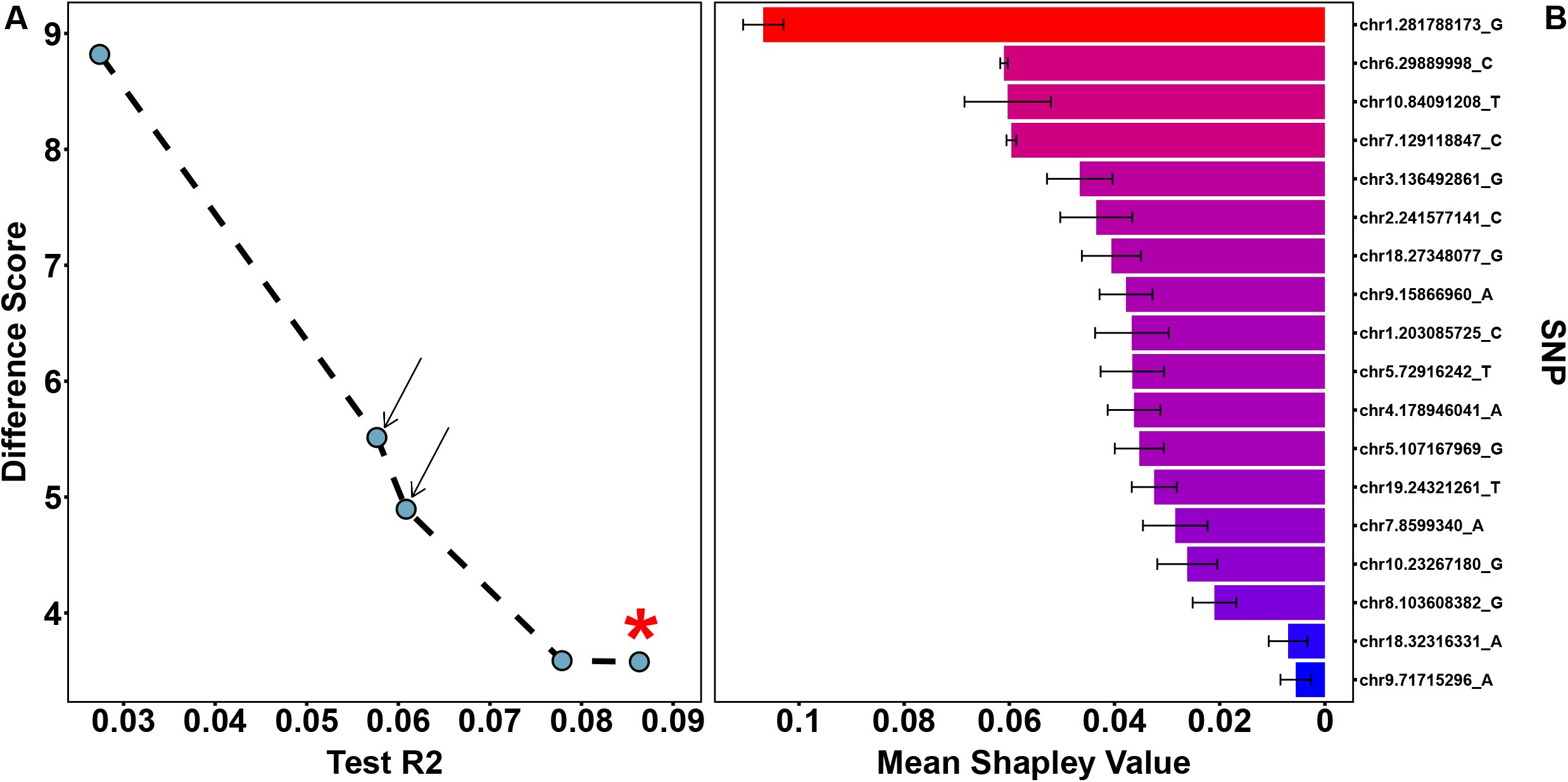
A. Final Pareto front of an example AutoQTL run from 18 QTL dataset. Pipelines (blue dots) with arrows are those that are optimized for both scoring metrics. The pipeline marked with a star is the pipeline with a test R^2^ matching the test R^2^ before GP was executed. B. Mean SHAP feature importance scores across the five pipelines from the example AutoQTL run from 18 QTL dataset for each locus. Black error bars represent S.E.M. Red-blue gradient denotes higher (red) and lower (blue) mean feature importance scores.

**Table 1.** Final Pareto optimal pipelines from an example run of the 18 QTL dataset (also depicted in Figure 2A). Pipelines in bold correspond to pipelines with arrows in Figure 2A which are optimized in both test R^2^ and DS (Pareto efficient). HP = Hyperparameter. The first pipeline (row), with * in Regressor column (LR), corresponds to the pipeline with a star in Figure 2A.

| 1<sup>st</sup> Operator (HP) | 2<sup>nd</sup> Operator (HP) | 3<sup>rd</sup> Operator (HP) | Regressor | $R^2$ | DS |
| --- | --- | --- | --- | --- | --- |
| - | - | - | LR* | 0.086 | 3.58 |
| - | - | - | LR | 0.078 | 3.59 |
| **Recessive Encoder** | **VT (0.05)** | - | **LR** | **0.061** | **4.90** |
| **VT (0.05)** | **VT (0.35)** | **Recessive** | **LR** | **0.058** | **5.51** |
| Recessive Encoder | Overdominance Encoder | GF (0.30) | LR | 0.027 | 8.82 |

The locus with the highest Mean Shapley value across pipelines in the example run’s final Pareto front is chr1.281788173_G, almost double that of the next highest locus (chr6.29889998; Figure 2B; File 5). This was also the SNP with the highest signal in the original GWAS (File S1). Three of the SNPs with the top five Shapley values also have the top five GWAS signals in the original study. However, overall, the rank order of SNPs in this analysis and the original GWAS are different. For instance, chr6.29889998 had the second highest average Shapley value but the second lowest GWAS signal. This is not surprising as Shapley feature importance values indicate predictive power of each feature (locus) regarding the model (pipeline) being tested and not a specific hypothesis test. We explore the differences between Shapley values and *p*-values further in the discussion.

Regarding GP pipeline evolution for the example run, only slight improvement of DS was observed over the first two GP generations while mean R^2^ reduces slightly in generation two before stabilizing in generation three and beyond (File S1). This is because some Pareto optimal pipelines with lower R^2^, but high DS, were not discovered via GP until later generations. Additionally, the 18 QTL dataset reaches optimization before 25 generations had elapsed. The mean optimization time for the 18 QTL dataset across the 10 runs occurred at 5.2 generations (File S5). This is likely due to the strong additive main effects of the dataset indicated by the low pipeline and root regressor diversity found in final Pareto fronts (mean of seven pipelines and 88.6% LR regressors; File S5). Additionally, larger datasets will likely yield more optimization potential.

### Patterns of Regressor Selection Emerge with Simulated Epistasis

With nine XOR interactions, AutoQTL selects machine learning regressors more frequently than LR. This was observed in all runs (25.3 machine learning regressors (89.1% (79.9% RF and 9.2% DT)) compared to 3.1 LR (10.9%) in final Pareto front on average; example in Figure 3A; Figure 3B; File S5). The final Pareto front of AutoQTL’s analysis of the XOR dataset yields higher pipeline diversity compared to LR (28.4 pipelines on average compared to seven in 18 QTL dataset; see Figure 3A for example Pareto front; File S5). Table 2 contains the final Pareto optimal pipelines selected in an example run of AutoQTL with the XOR dataset (AutoQTL random seed = 12). Pipelines shown in bold are Pareto efficient (optimized for both metrics) and have arrows pointing to them in Figure 3A. Across all ten runs with the XOR dataset, most pipelines had 2-level encoders as the final inheritance model (mean = 87.68%) followed by 3-level encoders (mean = 10.56%) and no encoders (e.g., additive model; mean = 1.76%) (Figure 3C; File S5).

**Figure 3.**
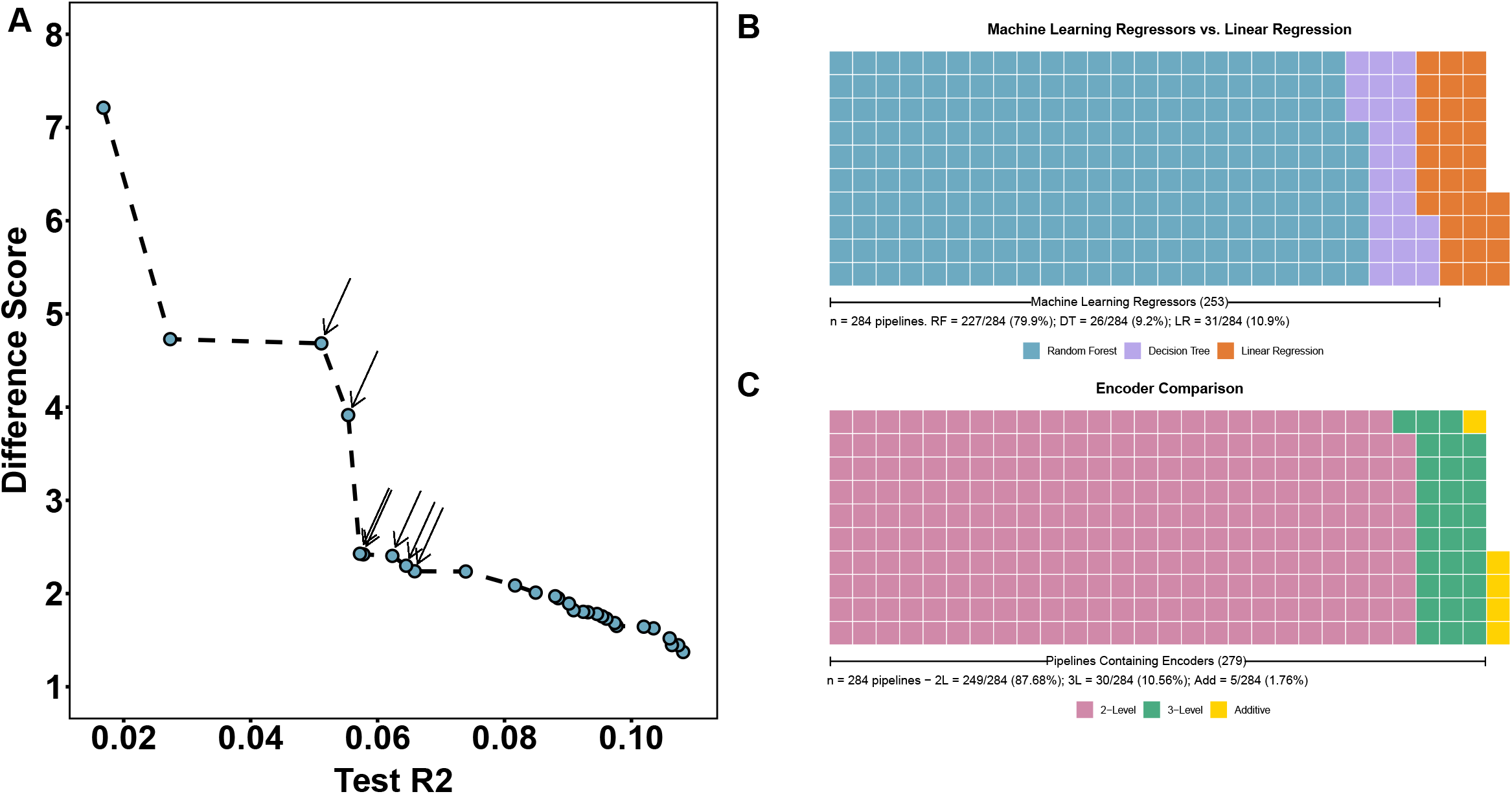
A. Final Pareto front of an example AutoQTL run from XOR (9 interaction) dataset. Pipelines (blue dots) with arrows are those that are optimized for both scoring metrics. B. Waffle plot of final Pareto front root regressor diversity across 10 AutoQTL runs of the XOR dataset (n = 284 total Pareto optimal pipelines). Each square of the plot represents one Pareto optimal pipeline. C. Waffle plot of final encoding state of loci across the same 10 runs (284 pipelines) in B.

**Table 2.** Final Pareto optimal pipelines from an example run of the XOR dataset (also depicted in Figure 3A). Pipelines in bold correspond to pipelines with arrows in Figure 3A which are optimized in both test R^2^ and DS (Pareto efficient). HP = Hyperparameter. Random Forest hyperparameters = bootstrap (T/F), max features, minimum samples per leaf, minimum samples per split. Decision Tree hyperparameters = max depth, minimum samples per leaf, minimum samples per split.

| 1<sup>st</sup> Operator (HP) | 2<sup>nd</sup> Operator (HP) | 3<sup>rd</sup> Operator (HP) | 4<sup>th</sup> Operator (HP) | 5<sup>th</sup> Operator (HP) | Regressor (HP) | $R^2$ | DS |
| --- | --- | --- | --- | --- | --- | --- | --- |
| Heterosis | - | - | - | - | RF (F, 0.2, 3, 12) | 0.108 | 1.37 |
| Heterosis | GF (0.1) | - | - | - | RF (T, 0.25, 3, 11) | 0.107 | 1.45 |
| Heterosis | Overdominance | Heterosis | Recessive | Recessive | RF (T, 0.25, 3, 11) | 0.106 | 1.45 |
| Heterosis | - | - | - | - | RF (T, 0.2, 3, 12) | 0.106 | 1.52 |
| Heterosis | Heterosis | - | - | - | RF (T, 0.25, 1, 20) | 0.103 | 1.63 |
| Heterosis | Heterosis | Underdominance | - | - | RF (T, 0.25, 3, 20) | 0.102 | 1.64 |
| Heterosis | - | - | - | - | RF (T, 0.4, 11, 5) | 0.098 | 1.65 |
| Heterosis | - | - | - | - | RF (T, 0.65, 14, 20) | 0.097 | 1.68 |
| Overdominance | Dominant | - | - | - | RF (T, 0.2, 8, 12) | 0.096 | 1.73 |
| Heterosis | Underdominance | - | - | - | RF (T, 1.0, 19, 20) | 0.095 | 1.76 |
| Heterosis | - | - | - | - | RF (T, 0.75, 19, 17) | 0.095 | 1.78 |
| Heterosis | - | - | - | - | RF (T, 0.35, 14, 20) | 0.093 | 1.80 |
| Heterosis | Underdominance | - | - | - | RF (T, 0.65, 19, 20) | 0.092 | 1.80 |
| Overdominance | Dominant | Heterosis | Overdominance | - | RF (T, 0.6, 19, 12) | 0.091 | 1.82 |
| Heterosis | Heterosis | - | - | - | RF (T, 0.25, 14, 20) | 0.090 | 1.89 |
| Heterosis | Heterosis | - | - | - | RF (T, 0.25, 16, 20) | 0.088 | 1.95 |
| Heterosis | Heterosis | - | - | - | RF (T, 0.25, 17, 20) | 0.088 | 1.97 |
| Heterosis | Heterosis | - | - | - | RF (T, 0.25, 19, 11) | 0.085 | 2.01 |
| Heterosis | Heterosis | VT (0.0) | - | - | RF (T, 0.2, 18, 12) | 0.082 | 2.09 |
| Heterosis | Underdominance | - | - | - | RF (T, 0.15, 19, 20) | 0.074 | 2.24 |
| **Overdominance** | **Dominant** | - | - | - | RF (T, 0.1, 16, 17) | 0.066 | 2.24 |
| **Overdominance** | **Dominant** | **Overdominance** | **Dominant** | **GF (0.35)** | RF (T, 0.1, 16, 12) | 0.065 | 2.30 |
| **Overdominance** | **Dominant** | **Overdominance** | **GF (0.35)** | - | RF (T, 0.1, 19, 17) | 0.062 | 2.40 |
| Heterosis | - | - | - | - | RF (T, 0.05, 19, 20) | 0.058 | 2.42 |
| **Overdominance** | **Dominant** | **Heterosis** | - | - | RF (T, 0.1, 19, 17) | 0.057 | 2.43 |
| **SP (65)** | **VT (0.3)** | **SP (10)** | **Heterosis** | - | DT (3, 9, 15) | 0.055 | 3.91 |
| **SP (55)** | **VT (0.3)** | **SP (10)** | **Heterosis** | - | DT (4, 1, 14) | 0.051 | 4.68 |
| - | - | - | - | - | DT (2, 5, 8) | 0.027 | 4.73 |
| SP (55) | VT (0.15) | VT (0.3) | SP (65) | Heterosis | LR | 0.017 | 7.21 |

Mean test R^2^ evolves more steadily than in the 18 QTL dataset and for more generations (19.5 generations on average compared to 5.2; File S5). DS slightly decreases over GP generations (File S1). This is likely due to more machine learning regressors being selected over GP generations which results in pipelines optimized for test R^2^ (likely overfit) increasing in prevalence (depicted in the lower right portion of Figure 3A).

Although somewhat arbitrary as all loci were involved in XOR interactions, we plotted average Shapley feature importance values (as violin plots) of each locus in this dataset in File S1. We hypothesized that features involved in epistatic interactions will have similar feature importance scores. This was mostly the case except for interaction pairs 3, 8, and 9. Due to the random shuffling process, we likely created stronger main effects in some loci over others, leading to these discrepancies. Generally, however, most interaction pairs appear to have similar values of feature importance.

### Linear Regression fails to capture V_p_ as Interactions are Added

The model R^2^ from LR of the dataset with all main effects removed via shuffling was relatively close to zero (mean = −0.00027; File S5). With each successive addition of an XOR interaction, the test R^2^ using LR alone remained stable with increases or decreases resulting from minor changes in main effects due to genotype shuffling (−0.0020 – 0.0085, on average). However, after each XOR addition, mean test R^2^ of machine learning regressors rose steadily with increasing interactions (Figure 4A). Over the course of the experiment, the mean test R^2^ of machine learning regressors increased by 0.016 on average per additional XOR interaction with the highest mean R^2^ (0.14) observed at nine interactions. The average number of machine learning regressors in the final Pareto fronts increased as soon one interaction was added (at zero interactions, mean of 2.4 pipelines (67%); at one interaction, mean of 9.2 pipelines (83.5%); at nine interactions, mean of 22.2 pipelines (83.5%); File S5). However, once reaching an average of 83.5% of pipelines at one interaction, the proportion of machine learning regressors remained relatively stable up until nine interactions (Figure 4C). Additionally, the average number of pipelines in the final Pareto front increased as interactions were added (3.6 on average at zero interactions; 25.5 on average at nine interactions). Encoder diversity remained relatively stable across the experiment with 2-level encoders being the final inheritance pattern selected in most final Pareto pipelines (mean = 79.2%; Figure 4E; File S5). Average 2-level encoder prevalence rose slightly over the course of the experiment (69% at zero interactions; 81% at nine interactions; Figure 4E; File S5).

**Figure 4.**
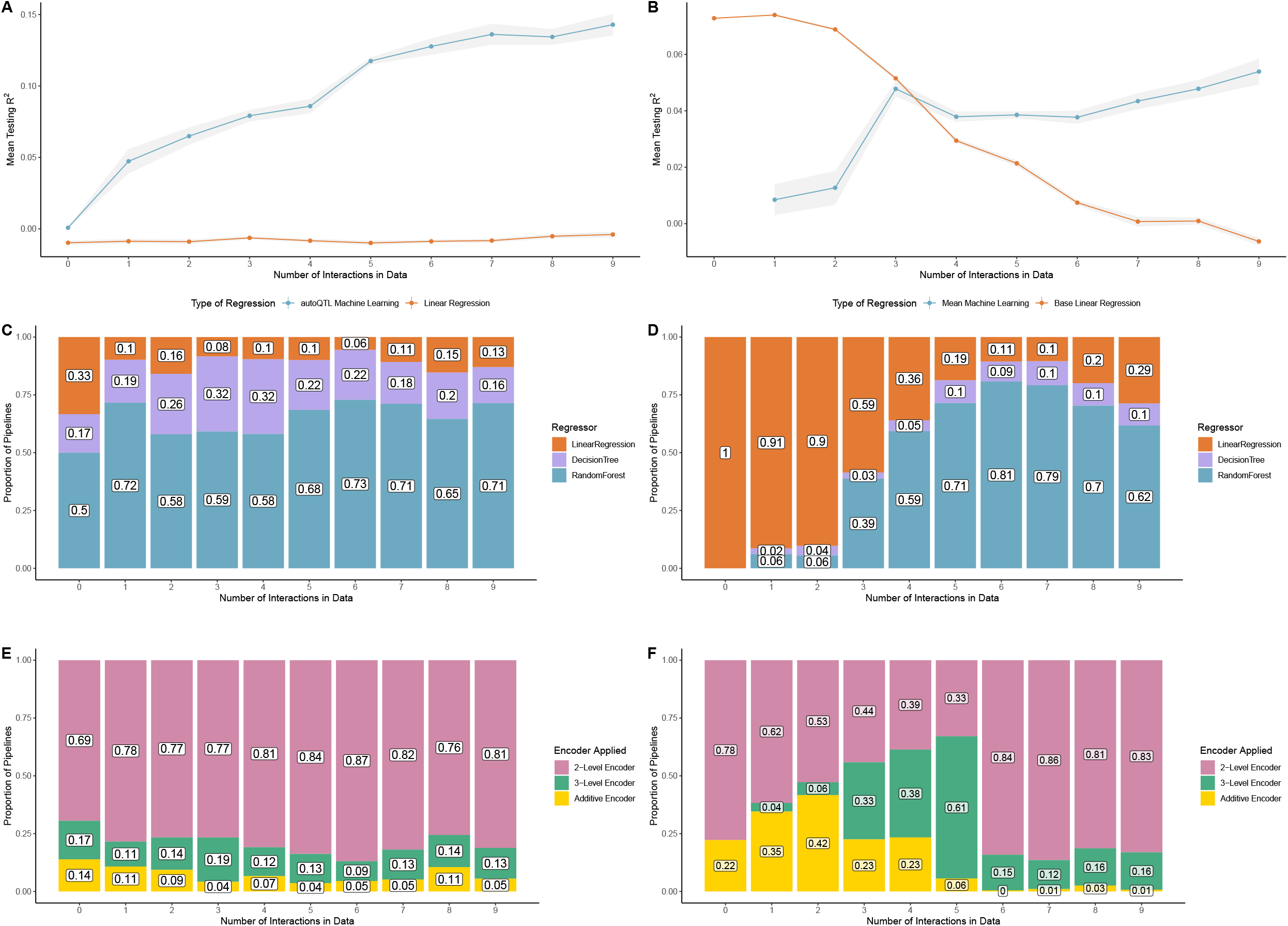
A. Mean test R2 of machine learning regression (DT and RF; blue dots and lines) and LR (orange dots and lines) for final Pareto optimal pipelines using dataset with random variables replaced by increasing XOR interactions. Gray shading around lines represents S.E. B. Mean test R^2^ of machine learning regression (DT and RF; blue dots and lines) and LR (orange dots and lines) for final Pareto optimal pipelines using datasets with putative QTL replaced by increasing XOR interactions. Gray shading around lines represents S.E. C. Stacked bar graphs illustrating the proportion of root regressors in final Pareto fronts with increasing number of epistatic pairs for datasets with random variables replaced by increasing XOR interactions. Orange bars = LR pipelines. Purple bars = DT pipelines. Blue bars = RF pipelines. Numbers inside bars represent respective proportions of each root regressor in that run. D. Stacked bar graphs illustrating proportion of root regressors in final Pareto fronts with increasing number of epistatic pairs for datasets with putative QTL replaced by increasing XOR interactions. Colors of bars and numbers in bars represent the same features as in C. E. Stacked bar graphs illustrating the proportion of encoder type in final Pareto fronts with increasing number of epistatic pairs for datasets with random variables replaced by increasing XOR interactions. Pink bars = 2-level encoders. Green bars = 3-level encoders. Yellow = no encoder selected (Additive encoding). Numbers inside bars represent respective proportions of encoder in that run. F. Stacked bar graphs illustrating the proportion of encoder type in final Pareto fronts with increasing number of epistatic pairs for datasets with increasing number of epistatic pairs for datasets with putative QTL replaced by XOR epistatic interactions. Colors of bars and numbers in bars represent the same features as in E.

Similar patterns were observed with the original 18 QTL dataset as main effects were replaced with interactions. With each successive removal of two loci with main effects and the addition of one XOR interaction, the mean test R^2^ using LR alone decreased from 0.0728 at zero interactions to −0.00629 at nine interactions while the mean test R^2^ of machine learning regressors increased from 0.00847 at zero intractions to 0.0539 at nine interactions (Figure 4B; File S5). Interestingly, there was a spike of average machine learning regressor test R^2^ at three interactions (0.0478). This value fell slightly (0.0378 at four interactions) until again increasing from six to nine interactions. Machine learning regressors replaced LR as the most prevalent root regressor in Pareto fronts at four interactions (average of 9.6 pipelines (62.3%)) and remained the most prevalent up to nine interactions (average of 19.4 pipelines (74.8%); Figure 4D: File S5). As observed in the previous experiment with random variables replaced by XOR interactions, pipeline number in the final Pareto front increased as interactions were added (8.4 on average at zero interactions; 28 on average at nine interactions; File S5). Unlike the previous experiment, encoder diversity shifted as interactions were added (Figure 4F). At zero interactions, most final encoders were 2-level (78% on average) while the remainder were additive (no encoder; 22% on average). This was due to many pipelines with lower test R^2^ and higher DS selecting two level encoders in the final Pareto front. However, it is important to note that all runs at zero interactions included pipelines with no encoders and high test R^2^ as seen in the example run in our 18 QTL experiment (pipeline with star in Figure 2A). As interactions increased from one to five, 3-level encoders rose in prevalence (mean of 4% at one interaction to a mean of 61% at five interactions). However, at six interactions and beyond, 2-level encoders made up most final inheritance models observed (83.5% on average from six to nine interactions).

The absolute value of the difference in Shapley feature importance in the example run for each interaction pair reduced for four pairs as loci transitioned from main effects to being involved in a 2-way XOR interaction (Figure 5A-D; File S5). However, there were two pairs where the difference increased (Pairs 5 and 9; File S5) while three pairs remained similar (Pairs 6, 7, and 8) (Figure 5E-I; File S5). These results are explored further in the discussion.

**Figure 5.**
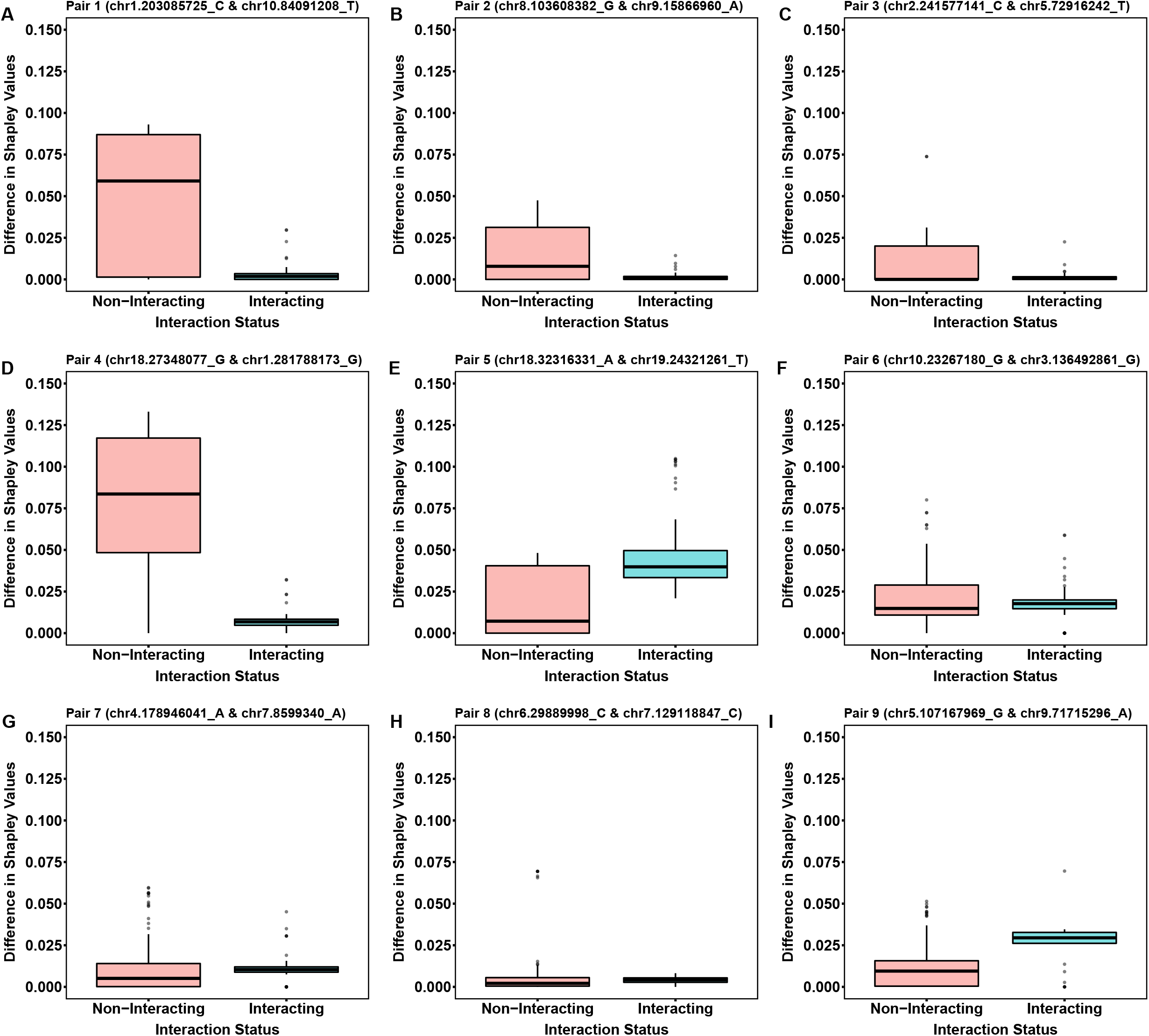
Boxplots illustrating the absolute value of the difference between Shapley feature importance values for loci while main effects are retained (Non-Interacting; red boxplots) and when they are part of an XOR interaction (Interacting; blue boxplots). Each boxplot (A-I) represents one of the nine possible interaction pairs. Locus names are in the title of each boxplot.

## Discussion

### Validation and Extensibility of AutoQTL

Results from the 18 QTL benchmark data validate AutoQTL’s ability to capture the total *V_P_* via a Pareto optimal pipeline. This is the pipeline with the highest test R^2^ but the lowest DS (Figure 2A, Table 1). In the example run illustrated in the results, there are four additional pipelines that explain less V_*P*_ but are less overfit (higher DS) and, thus, are more generalizable. This generalizability is observed by these pipelines’ performance on holdout data where the increase in R^2^ is proportionally higher than more overfit models (File S4). In addition to higher generalizability, these pipelines have additional operators selected, with most having features (putative QTLs) removed and some performing both feature selection and encoding, as in the two bolded pipelines in Table 1 from the example run. These additional pipelines may be identifying false positives from the original GWAS via removal of features that do not meet the criteria of the feature selection hyperparameters. Our feature selection methods employ two distinct strategies. The VT and GF feature encoders select features based upon metrics of allele/genotype diversity while SP selects features by employing f-regression and the association between feature and phenotype [21]. The latter strategy (SP) likely has a greater potential to sort out false positives. However, VT and GF, which were chosen in both bolded example pipelines in Table 1 and all additional pipelines in the example final Pareto front from the 18 QTL dataset (File S4), also have the potential to remove false positives as features with lower minor allele frequencies have inherently less heterozygosity and are usually less informative [38].

The addition of encoder operators is interesting in that there are likely QTL within the 18 QTL dataset with significant deviations from additivity. In other words, the V_*P*_ explained by these loci increase when converted to an inheritance model other than additive. This was observed for four loci (chr1.281788173_G – Recessive; chr8.103608382_G – Dominant; chr9.71715296_A – Underdominant; chr18.27348077_G – Recessive; File S1). Of these, chr1.281788173 also had the highest average Shapley feature importance values. This likely drove the selection of pipelines that removed other less predictive loci and loci that were not well described using 2-level encoders. This explains why so many 2-level encoder pipelines were found in our final experiment when no interactions were added. The predictive value of chr1.281788173 in its proper encoding (recessive) was likely so high that many Pareto optimal pipelines were built around this optimization strategy. Indeed, the average Shapley feature importance of this locus under an additive inheritance model is 0.089 while under the recessive model it is 0.118 (File S5). This is further illustrated by three of the five Pareto optimal pipelines in the example run of the 18 QTL dataset selecting the recessive encoder (Table 1; File S4) and 78% of Pareto optimal pipelines in our final experiment having 2-level encoders at zero interactions (Figure 4F). These results illustrate that AutoQTL has the potential to identify deviations from additivity via feature selection and encoding operators. In future versions, we plan on adding encoding checks into both feature importance strategies and in pipeline construction to identify deviations from additivity on a locus-by-locus basis.

Shapley value-based Feature importance values place SNPs in a different rank order than the *p*-value rank order in the GWAS results (File S1). This is not surprising for multiple reasons. The first is that the original GWAS p-values were derived from millions of independent tests while our feature importance values are only from a total of 18 features. More importantly though, feature importance metrics are not linked to hypothesis testing. Rather, feature importance metrics indicate the predictive power of each feature in their respective model (pipeline in the case of AutoQTL). Another reason these ranks are different is because AutoQTL provides the user multiple Pareto optimal pipelines, each with different models potentially containing feature selection and/or encoding phases. As we have discussed, encoding changes can alter each feature’s importance, sometimes drastically. Additionally, if a feature is removed from a pipeline, its feature importance is zero, considerably affecting the mean in most cases. Due to the combined effects of potentially variable feature importance values across models and the dissimilar interpretations of feature importance metrics compared to *p*-values, researchers should compare the results of AutoQTL and GWAS/QTL analysis with caution as both approaches are asking different questions. However, what feature importance metrics offer are new and/or alternative insights into genotype/phenotype associations and putative variant identification. This, coupled with the diverse pipelines produced by AutoQTL, can lead researchers to new discoveries in quantitative genetics.

### Evidence for Epistasis Detection in AutoQTL

In the presence of nine XOR interactions, AutoQTL’s final Pareto front is populated by more pipelines selecting machine learning regression (25.3 (89.1%) machine learning vs. 3.1 LR (10.9%) on average; Figure 3B; File S5; File S6) compared to final Pareto front of the 18 QTL dataset (0.8 (11.4%) machine learning vs. 6.2 LR (88.6%) on average; File S4). The XOR dataset also optimizes over more generations (19.5 in XOR vs. 5.2 in 18 QTL on average; File S5; File S7) and has higher count of final Pareto optimal pipelines compared to the LR dataset analysis (28.4 in XOR vs. 7 in 18 QTL on average; File S5).

The proportion of machine learning regressors selected in the XOR dataset is partially attributable to the ability of machine learning regressors (especially RF) to detect interactions [6]. Indeed, our results show that AutoQTL has the potential to detect epistatic interactions. However, the longer optimization period and greater pipeline count/regressor diversity observed in the XOR analysis has two other possible explanations. The first is that machine learning regressors have a higher tendency to overfit compared to LR, leading to potentially more Pareto optimal solutions being generated (as seen in the lower right portion of Figure 3A). The second, and likely most considerable, is that both RF and DT have several hyperparameters that can be selected by AutoQTL whereas LR has none, expanding both the nodes that can be modified by future rounds of selection and, therefore, the number of possible pipelines that can be generated. However, these diverse pipelines can yield multiple points of evidence for the strength of certain interactions over others or compare the strength of interactions to main effects in real world data. Additionally, diverse pipelines offer more model options to researchers for future inquiry.

The vast majority of Pareto optimal pipelines in the nine XOR analysis had at least one feature encoder selected (98.2% of pipelines; Figure 3C; File S5). Of the 279 pipelines that selected encoders, 249 resulted in a 2-level final encoding. That is, two genotypic states instead of three. In the 18 QTL dataset and likely all datasets with only main effects, 2-level encoding is likely due to large effect QTL better explained with dominance deviations or heterosis. However, in a dataset made up of 9 XOR interactions, as in experiment 2, another possible explanation could be that 2-level encodings better match the strict penetrance function of XOR, in which two-locus genotypes result in a penetrance of 1 or 0. It may be possible that AutoQTL is attempting to restructure the data in a format that better mirrors the complete penetrance of the XOR model. More testing with other types of interactions, including those that model incomplete penetrance and cartesian products, is needed to verify this. Since there are potentially four values a cartesian product can take with standard additive encodings (0, 1, 2, and 4), 2-level encoding prevalence may only be a detectable signal of epistasis in full penetrance models or in epistatic interactions involving one or more non-additive loci.

In the experiment where one XOR interaction was added at a time to a dataset of little to no main effects, we observed that the R^2^ detectable my machine learning regressors increased with successive interactions while R^2^ detectable by LR remained low (Figure 4A; File S5; File S8). This provides further evidence that machine learning regressors have the capability detect epistatic interactions. We also observed that even as low as one added interaction, the proportion of machine learning regressors in the final pareto front increased by approximately 24% on average (Figure 4C; File S5; File S8). However, the change was not as stark as in the final experiment when interactions were added to main effects. After this, the relative proportion of regressors remained mostly unchanged from one to nine total interactions. This is likely due to machine learning regressors able to describe more variance when there are very low main effects compared to LR and the propensity of machine learning regressors to overfit. A similar result was observed with encoders as the overall proportion of encoder types changed little across the experiment (Figure 4E; File S5; File S8). However, what this experiment importantly illustrates is that AutoQTL can detect increasing levels of epistatic signal as interactions increase in a dataset.

### Epistatic Detection and Feature Importance in the Presence of Main Effects

Like the experiment where random features are replaced by XOR interactions, we observed that as main effects are removed and converted to XOR interactions, the R^2^ detectable by machine learning regressors increases. However, along with this comes a concurrent reduction in R^2^ detectable by LR (Figure 4B; File S5; File S9). This was expected but provides further evidence of the ability of machine learning regressors (particularly RF) to detect epistasis over LR. In contrast to the previous experiment, however, we observed strong trends in regressor and encoder diversities as main effects were replaced by interactions. At zero interactions, the only regressor to population the final Pareto front was LR due to the strong main effects (Figure 4D; File S5; File S9). However, at three total interactions, the proportion of machine learning regressors began to increase and overtook LR as four interactions were added. The proportion of machine learning regressors remained high up to nine total interactions. This illustrates that machine learning regressors, driven by RF, can more accurately describe epistatic relationships over LR. However, along with this trend also comes the inflation of the final Pareto front with more optimal pipelines (File S5). As with previous experiments, this is likely due to these pipelines having a higher potential to overfit as well as having more hyperparameters to optimize. The proportion of 3-level encoders increased with each successive interaction added from one to five total interactions before falling sharply at six total interactions (Figure 4F). It is unclear why this occurred. One explanation is, because this pattern occurred when there were similar instances of 2-way interactions and single loci with large main effects, this type of encoder pattern arises when there is a mix of main effects and epistatic signals in a dataset. Further analysis is required to strengthen this explanation as this, if true, may only occur with interactions modeled with pure penetrance as in XOR. However, when main effects reduced in number, 2-level encoders became much more prevalent which is what was also observed in the datasets with nine XOR interactions and in the experiment with increasing XOR interactions replacing random features. In future experiments, we aim to model features that have strong main effects and are also involved in epistatic interactions with other loci. It has been observed that loci with strong main effects are also epistatic hubs, involved in many 2- and 3-way epistatic interactions in the genome [3]. We hope to discover these epistatic hubs as well as epistatic loci without large main effects in future experiments using AutoQTL.

Figure 5 illustrated that for four pairs of loci, the absolute difference in feature importance values reduce as loci lose their main effects and become part of a 2-way interaction. However, for the remaining five pairs, the difference in feature importance either increases or remains similar. We originally hypothesized that feature importance scores of interacting loci would converge as they lost their main effects and become part of an interaction pair. However, there are two main reasons why this wasn’t observed universally. The first is that these are manufactured interactions with similar proportions of the genotype scores being used to generate the interaction. This leads to similar signals and therefore, likely very similar feature importance metrics. The second is that some loci already exhibit similar feature importance scores with main effects (Figure 2B; File S5). The act of removing the main effects of each locus and generating an interaction signal led to a greater difference in feature importance scores for some pairs, perhaps by chance. It is our goal in future experiments to detect true epistatic relationships and compare feature importance metrics of additive vs. non-additive effect loci.

### Limitations and Future Expansion of AutoQTL

One potential criticism of the approach, likely to arise from trained geneticists, is that AutoQTL provides too many options (i.e., Pareto optimal predictive models). How should one approach selecting which model to apply to their data? This is a valid concern that deserves a focused response. In addition to understanding that prediction is wholly distinct from hypothesis testing, machine learning approaches have the potential to overfit the training set and not become generalizable to other datasets. We chose to address this issue by invoking a Pareto optimization strategy which employs two optimization metrics with one specifically tailored to address overfitting (DS). Furthermore, the graph-based expression tree structure of our pipelines allows us to explore more dimensions than standard approaches can including non-additive effects, feature selection, regressor type, and hyperparameter optimization. Due to these factors, there are potentially many equally optimal predictive models for any dataset. However, we recommend researchers think about Pareto efficiency when selecting a pipeline (model). That is, ones that are optimized in both scoring metrics like the pipelines we have outlined with arrows in Figures 2A and 3A and in bold in Tables 1 and 2. Furthermore, we recommend running AutoQTL multiple times with different random seeds (different training/testing data splits) to allow the GP to explore and discover new pipelines. Even though AutoQTL is still able to explain all the V_*p*_ in the standard linear model (18 QTL dataset experiment), the Pareto efficient pipelines tell us important information about loci that have strong non-additive effects and loci that could be false positives in the GWAS. Thus, AutoQTL has the capability to elucidate genetic relationships that are extremely difficult or impossible to reveal using standard approaches.

As are many autoML methods to maintain computational tractability, AutoQTL is currently limited in the number of features that can be analyzed through its automated workflow. One major benefit of standard QTL analysis is that there is no theoretical limit to the number of loci that can be analyzed as QTL analysis and GWAS employ modified versions of univariate regression. However, AutoQTL uses multiple LR as a base regressor option and can therefore not have more features than observations. However, we are currently developing ways to address this limitation by developing pruning strategies that identify important features from large GWAS-level datasets using autoML methods, statistical thresholds, and/or *a priori* expert biological knowledge.

We have currently added Shapley values as an AutoQTL feature importance module. However, in the future, we wish to modify our current feature importance strategy to consider both univariate effects and epistatic interactions so that researchers are more informed of the genetic variance underlying important traits of interest and identify undiscovered genetic relationships. Furthermore, we wish to apply encoders on a locus-by-locus basis to produce genotype values indicative of the actual inheritance model of each SNP to better describe each feature’s importance to the model being tested.

In addition to the encoder and feature selection operator classes we currently employ, we aim to add other classes of operators that can contain detailed structural and function genomic information. Examples of this would be if a SNP is located in a gene model or regulatory region, if the gene(s) associated with a particular SNP is part of a certain biological pathway or biological function, and if there is expert-based knowledge on a SNP’s effect on phenotypes, genes, and/or other biological entities. Operator classes like this will allow AutoQTL to be more informed about each SNP which has the potential to enhance downstream analyses as well as putative QTL detection.

Finally, we are currently developing a feature which will provide researchers with executable Python code of the models generated in AutoQTL’s final Pareto front. This will allow easy application of AutoQTL models to genomic data for further analysis.

## Conclusions

We have demonstrated that AutoQTL can both capture V_*P*_ in a dataset of putative QTL from real-world experimentation and provide evidence of epistatic interactions in simulated data derived from real-world data. In addition, our optimization strategy provides diverse solutions that differ from standard QTL analysis using a set of feature selectors, feature encoders, and machine learning regressors with a varied set of hyperparameters. Feature importance adds to this by providing researchers a different perspective on selecting potential targets of further study compared to standard statistical approaches. We intend to further improve the software to both enrich and supplement the analysis of complex traits in both human and non-human systems. It is our aim that AutoQTL can assist in the spaces of precision medicine and quantitative trait analysis by enhancing the exploration of diverse phenotypes, disease models, and non-additive effects in all omics-level datasets.

AutoQTL is open-source and available on Github: https://github.com/EpistasisLab/autoqtl.

## List of Abbreviations

DEAP: Distributed Evolutionary Algorithms in Python
DS: Difference Score
DT: Decision Tree (regressor)
GF: Genotype Frequency (feature selector)
GP: Genetic Programming
GWAS: Genome-Wide Association Study
LR: Linear Regression (regressor)
NSGA: Non-Dominated Sorted Genetic Algorithm II
PCA: Principal Component Analysis
QTL: Quantitative Trait Locus
RF: Random Forest (regressor)
SHAP: Shapley Additive exPlanations
SNP: Single Nucleotide Polymorphism
SP: Select Percentile (feature selector)
TPOT: Tree-based Pipeline Optimization Tool
V_*P*_: Phenotypic Variance
VT: Variance Threshold (feature selector)

## Declarations

### Ethics approval and consent to participate

Not applicable.

### Consent for publication

Not applicable.

### Availability of data and materials

Raw data to run all experiments are found in File S10. File S2 contains R scripts to generate datasets with varying numbers of XOR interaction pairs.

### Authors’ contributions

Manuscript was written by PJF and AG. PJF analyzed data and generated figures and tables. PJF, RL, and JHM outlined experiments. AG developed the software, ran experiments 1 and 2, and organized data. EZ and TL developed genotype shuffling techniques and ran experiments 3 and 4. RL and JHM provided guidance and ideas concerning the scope of the software and manuscript and provided feedback on the manuscript. AC and AAP assisted with statistical analysis and GWAS data availability. All other authors, including AC and AAP, performed the original GWAS analysis and provided feedback on the manuscript.

## Acknowledgements

Not applicable.

## References

1. Miles CM, Wayne M. Quantitative Trait Locus (QTL) Analysis. Nature Education. 2008;1:208.

2. Wei W-H, Hemani G, Haley CS. Detecting epistasis in human complex traits. Nat Rev Genet. Nature Publishing Group; 2014;15:722–33.

3. Matsui T, Mullis MN, Roy KR, Hale JJ, Schell R, Levy SF, et al. The interplay of additivity, dominance, and epistasis on fitness in a diploid yeast cross. Nat Commun. Nature Publishing Group; 2022;13:1463.

4. Hallin J, Märtens K, Young AI, Zackrisson M, Salinas F, Parts L, et al. Powerful decomposition of complex traits in a diploid model. Nat Commun. Nature Publishing Group; 2016;7:13311.

5. Adams SM, Feroze H, Nguyen T, Eum S, Cornelio C, Harralson AF. Genome Wide Epistasis Study of On-Statin Cardiovascular Events with Iterative Feature Reduction and Selection. Journal of Personalized Medicine. Multidisciplinary Digital Publishing Institute; 2020;10:212.

6. Orlenko A, Moore JH. A comparison of methods for interpreting random forest models of genetic association in the presence of non-additive interactions. BioData Mining. 2021;14:9.

7. Ritchie MD, Hahn LW, Roodi N, Bailey LR, Dupont WD, Parl FF, et al. Multifactor-Dimensionality Reduction Reveals High-Order Interactions among Estrogen-Metabolism Genes in Sporadic Breast Cancer. The American Journal of Human Genetics. 2001;69:138–47.

8. Gelfman S, Wang Q, McSweeney KM, Ren Z, La Carpia F, Halvorsen M, et al. Annotating pathogenic non-coding variants in genic regions. Nat Commun. Nature Publishing Group; 2017;8:236.

9. Ioannidis NM, Rothstein JH, Pejaver V, Middha S, McDonnell SK, Baheti S, et al. REVEL: An Ensemble Method for Predicting the Pathogenicity of Rare Missense Variants. The American Journal of Human Genetics. 2016;99:877–85.

10. Olson RS, Bartley N, Urbanowicz RJ, Moore JH. Evaluation of a Tree-based Pipeline Optimization Tool for Automating Data Science. Proceedings of the Genetic and Evolutionary Computation Conference 2016 [Internet]. New York, NY, USA: Association for Computing Machinery; 2016 [cited 2022 Jul 18]. p. 485–92. Available from: https://doi.org/10.1145/2908812.2908918

11. Olson RS, Urbanowicz RJ, Andrews PC, Lavender NA, Kidd LC, Moore JH. Automating Biomedical Data Science Through Tree-Based Pipeline Optimization. In: Squillero G, Burelli P, editors. Applications of Evolutionary Computation. Cham: Springer International Publishing; 2016. p. 123–37.

12. Le TT, Fu W, Moore JH. Scaling tree-based automated machine learning to biomedical big data with a feature set selector. Bioinformatics. 2020;36:250–6.

13. Manduchi E, Fu W, Romano JD, Ruberto S, Moore JH. Embedding covariate adjustments in tree-based automated machine learning for biomedical big data analyses. BMC Bioinformatics. 2020;21:430.

14. Orlenko A, Moore JH, Orzechowski P, Olson RS, Cairns J, Caraballo PJ, et al. Considerations for automated machine learning in clinical metabolic profiling: Altered homocysteine plasma concentration associated with metformin exposure. Pac Symp Biocomput. 2018;23:460–71.

15. Orlenko A, Kofink D, Lyytikäinen L-P, Nikus K, Mishra P, Kuukasjärvi P, et al. Model selection for metabolomics: predicting diagnosis of coronary artery disease using automated machine learning. Bioinformatics. 2020;36:1772–8.

16. Manduchi E, Le TT, Fu W, Moore JH. Genetic Analysis of Coronary Artery Disease Using Tree-Based Automated Machine Learning Informed By Biology-Based Feature Selection. IEEE/ACM Transactions on Computational Biology and Bioinformatics. 2022;19:1379–86.

17. Langdon WB, Poli R, McPhee NF, Koza JR. Genetic Programming: An Introduction and Tutorial, with a Survey of Techniques and Applications. In: Fulcher J, Jain LC, editors. Computational Intelligence: A Compendium [Internet]. Berlin, Heidelberg: Springer; 2008 [cited 2022 Jul 18]. p. 927–1028. Available from: https://doi.org/10.1007/978-3-540-78293-3_22

18. Banzhaf W, Francone FD, Keller RE, Nordin P. Genetic programming: an introduction: on the automatic evolution of computer programs and its applications. San Francisco, CA, USA: Morgan Kaufmann Publishers Inc.; 1998.

19. Koza JR. Genetic Programming: On the Programming of Computers by Means of Natural Selection. Cambridge, MA, USA: Bradford Books; 1992.

20. Van Rossum G, Drake FL. Python 3 Reference Manual. Scotts Valley, CA: CreateSpace; 2009.

21. Pedregosa F, Varoquaux G, Gramfort A, Michel V, Thirion B, Grisel O, et al. Scikit-learn: Machine Learning in Python. Journal of Machine Learning Research. 2011;12:2825–30.

22. Doolittle DP. Dominance Deviations. In: Doolittle DP, editor. Population Genetics: Basic Principles [Internet]. Berlin, Heidelberg: Springer; 1987 [cited 2022 Jul 18]. p. 164–8. Available from: https://doi.org/10.1007/978-3-642-71734-5_36

23. Cormen TH, Leiserson CE, Rivest RL, Stein C. Introduction to Algorithms, Second Edition. 2nd edition. Cambridge, Mass: The MIT Press; 2001.

24. Jin Y. Multi-Objective Machine Learning. Berlin, Germany: Springer Science & Business Media; 2006.

25. Olson RS, Urbanowicz RJ, Andrews PC, Lavender NA, Kidd LC, Moore JH. Automating Biomedical Data Science Through Tree-Based Pipeline Optimization. In: Squillero G, Burelli P, editors. Applications of Evolutionary Computation. Cham: Springer International Publishing; 2016. p. 123–37.

26. Fortin F, De Rainville F, Gardner M, Parizeau M, Gagné C. DEAP: evolutionary algorithms made easy. J Mach Learn Res. 2012;13:2171–5.

27. Deb K, Pratap A, Agarwal S, Meyarivan T. A fast and elitist multiobjective genetic algorithm: NSGA-II. IEEE Transactions on Evolutionary Computation. 2002;6:182–97.

28. Lundberg SM, Lee S-I. A Unified Approach to Interpreting Model Predictions. Advances in Neural Information Processing Systems [Internet]. Curran Associates, Inc.; 2017 [cited 2022 Oct 22]. Available from: https://proceedings.neurips.cc/paper/2017/hash/8a20a8621978632d76c43dfd28b67767-Abstract.html

29. Chitre AS, Polesskaya O, Holl K, Gao J, Cheng R, Bimschleger H, et al. Genome-Wide Association Study in 3,173 Outbred Rats Identifies Multiple Loci for Body Weight, Adiposity, and Fasting Glucose. Obesity. 2020;28:1964–73.

30. Chitre AS, Polesskaya O, Holl K, Gao J, Cheng R, Bimschleger H, et al. Genome-Wide Association Study in 3,173 Outbred Rats for Body Weight, Adiposity, and Fasting Glucose [Internet]. Genes and Addiction: NIDA Center for GWAS in Outbred Rats. 2022 [cited 2022 Jul 18]. Available from: https://cgord.org/dataset/2

31. Chang CC, Chow CC, Tellier LC, Vattikuti S, Purcell SM, Lee JJ. Second-generation PLINK: rising to the challenge of larger and richer datasets. GigaScience. 2015;4:s13742-015-0047–8.

32. R Core Team. R: A language and environment for statistical computing [Internet]. Vienna, Austria: R Foundation for Statistical Computing; 2022. Available from: https://www.R-project.org/

33. Bateson W, Mendel G, Leighton AG. Mendel’s principles of heredity, by W. Bateson [Internet]. Cambridge, UK: Cambridge University Press; 1909. p. 1–448. Available from: https://www.biodiversitylibrary.org/bibliography/1057

34. Li W, Reich J. A Complete Enumeration and Classification of Two-Locus Disease Models. HHE. Karger Publishers; 2000;50:334–49.

35. Urbanowicz RJ, Kiralis J, Sinnott-Armstrong NA, Heberling T, Fisher JM, Moore JH. GAMETES: a fast, direct algorithm for generating pure, strict, epistatic models with random architectures. BioData Min. 2012;5:16.

36. Hastie T, Tibshirani R, Friedman J. The Elements of Statistical Learning: Data Mining, Inference, and Prediction. 2nd edition. New York, NY: Springer; 2016.

37. McKinney BA, Reif DM, Ritchie MD, Moore JH. Machine Learning for Detecting Gene-Gene Interactions. Appl-Bioinformatics. 2006;5:77–88.

38. Botstein D, White RL, Skolnick M, Davis RW. Construction of a genetic linkage map in man using restriction fragment length polymorphisms. Am J Hum Genet. 1980;32:314–31.

