## Supplementary material for "Automated quantitative trait locus analysis (AutoQTL)": FileS1

Supplementary File S1

I. Encoders

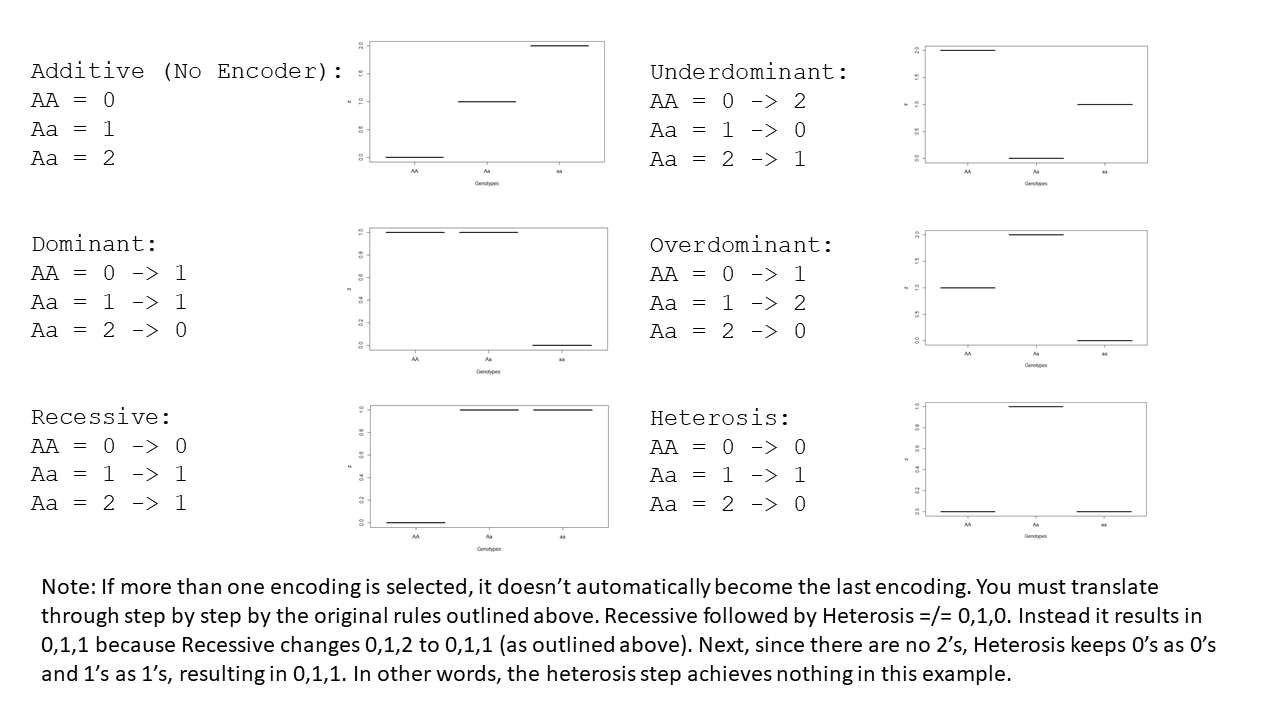

Note: If more than one encoding is selected, then a serial encoding occurs. It does not revert to additive and then apply the new encoder. Instead, genotypes are translated step by step by the original rules outlined in the images above. Below is an example where the Recessive Encoder is followed by Heterosis. In this example, the second encoding has no effect because there are no 2’s to change to 0’s.

```
Additive(Default) -> Recessive -> Heterosis
AA = 0            -> 0         -> 0
Aa = 1            -> 1         -> 1
Aa = 2            -> 1         -> 1
```

In this next example, the first and second encoders are both overdominant. However, the resulting inheritance model is Underdominant.

```
Additive(Default) -> Overdominant -> Overdominant
AA = 0            -> 1            -> 2
Aa = 1            -> 2            -> 0
Aa = 2            -> 0            -> 1
```

In the final example, two encoders lead to an inheritance model not possible with only one encoder. However, even though this numeric representation is not possible with one encoder, it is still recessive in terms of linear regression.

```
Additive(Default) -> Recessive    -> Overdominant
AA = 0            -> 0            -> 1
Aa = 1            -> 1            -> 2
Aa = 2            -> 1            -> 2
```

II. Example expression tree (autoQTL pipeline) with component parts identified

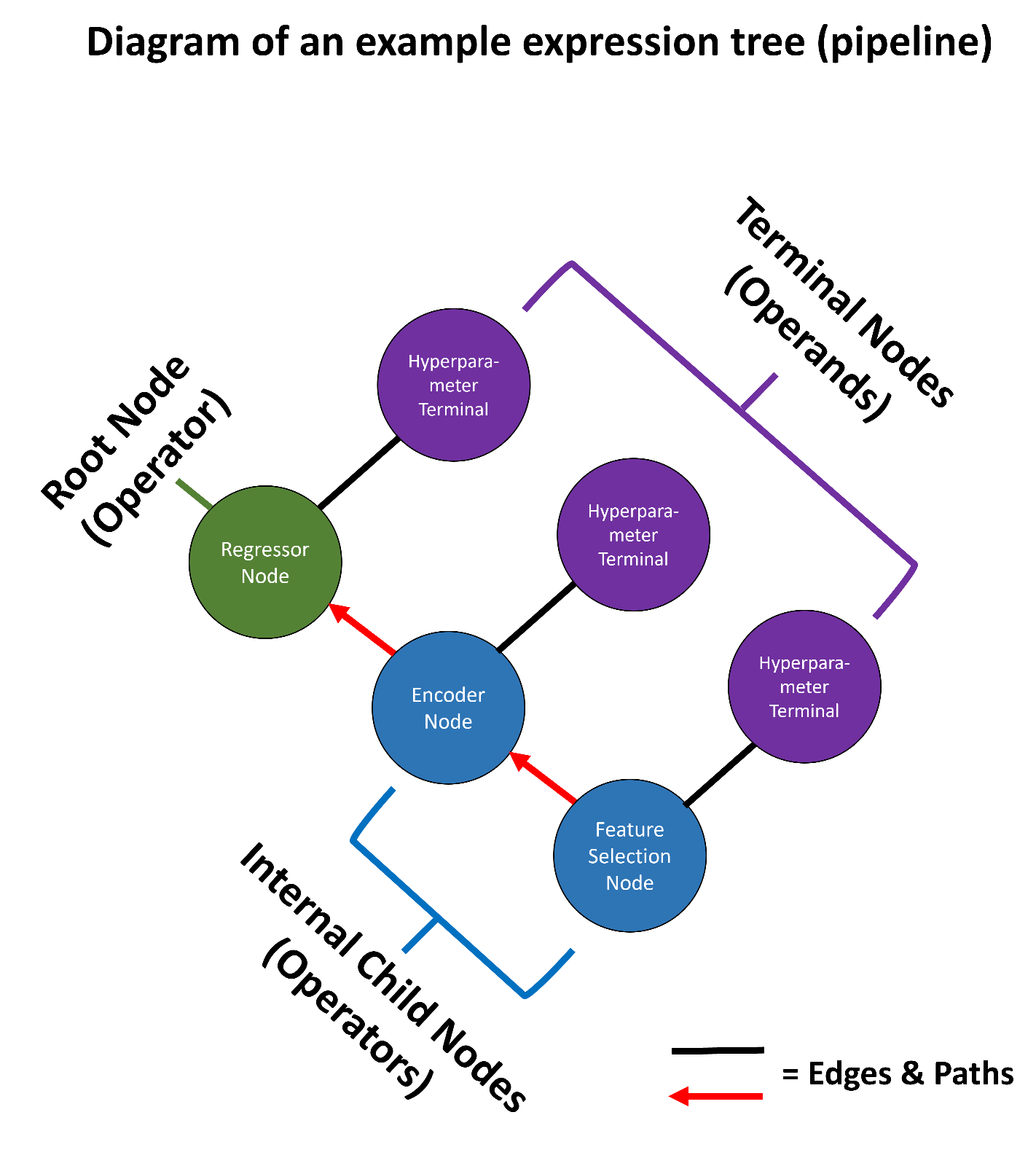

III. Mutation and Crossover Examples

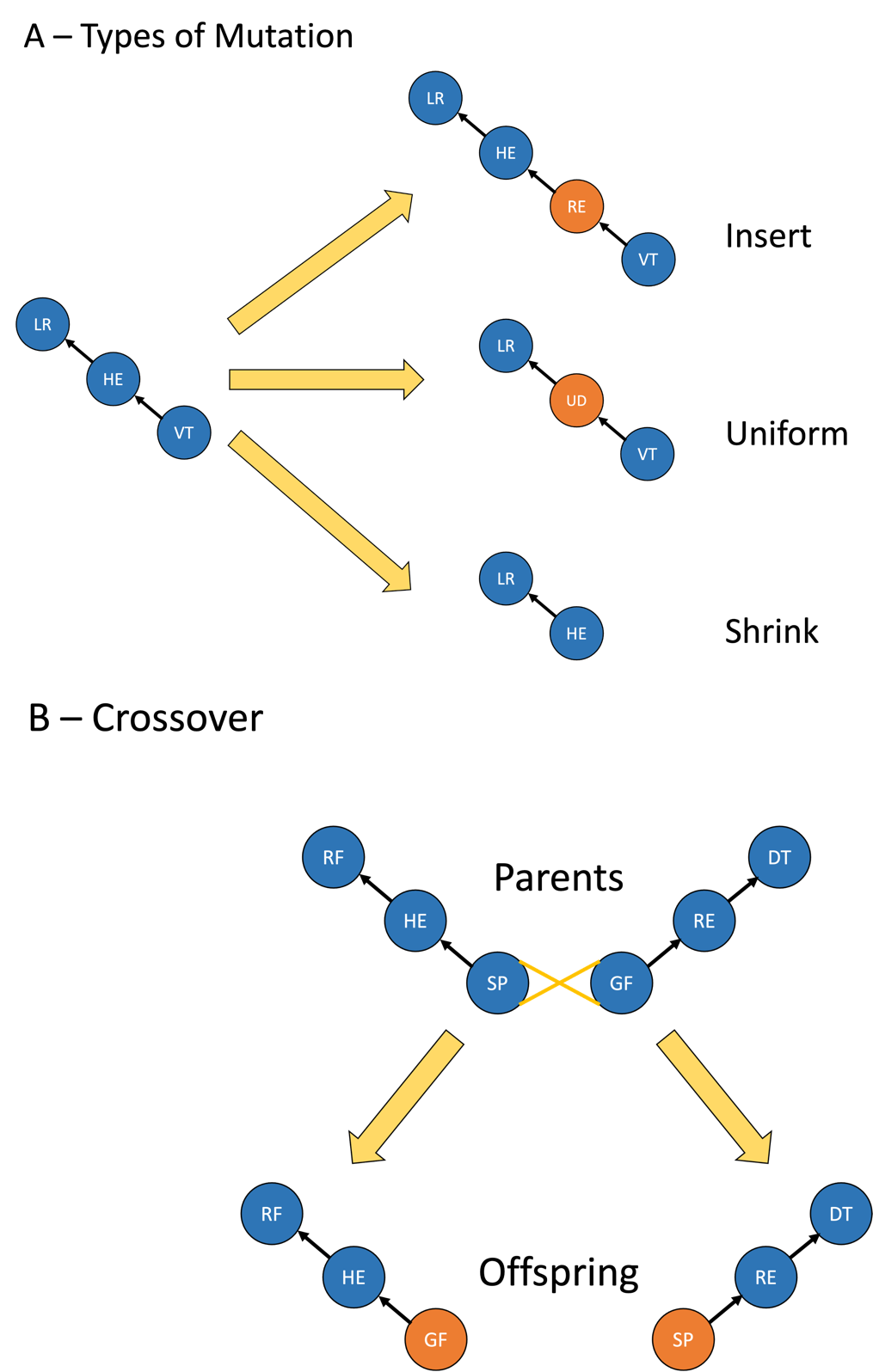

A. Types of mutation possible in AutoQTL with examples. B. An example crossover event in AutoQTL. The crossover occurred where the yellow “X” is shown. The operators are swapped to generate two offspring. Only the first offspring generated will be selected for the next generation. The other is discarded. Note mutation and crossover can also occur at hyperparameter (terminal) nodes (not shown). LR = Linear Regression, RF = Random Forest, DT = Decision Tree, HE = Heterosis Encoder, RE = Recessive Encoder, UD = Underdominance Encoder, VT = Variance Threshold Feature Selection, SP = Select Percentile Feature Selection, GF = Genotype Frequency Feature Selection

IV. XOR Penetrance Function

|  | **SNP 2** | | | | **Marginal penetrance** |
| --- | --- | --- | --- | --- | --- |
|  | **Genotype** | **BB (0)** | **Bb (1)** | **bb (2)** |  |
| **SNP 1** | **AA (0)** | 0 | 1 | 0 | 0.5 |
|  | **Aa (1)** | 1 | 0 | 1 | 0.5 |
|  | **aa (2)** | 0 | 1 | 0 | 0.5 |
|  | **Marginal Penetrance** | 0.5 | 0.5 | 0.5 | K = 0.5 |

The table above describes the XOR penetrance function. XOR is both a strict and pure penetrance function. Strict in that all the loci (in this case, 2) are predictive of the phenotype and pure in that the interaction does not display main effects [1]. XOR is a fully penetrant and purely epistatic model in which each two-locus genotype results in a marginal penetrance of exactly 0.5.

**References**

1. Urbanowicz RJ, Kiralis J, Sinnott-Armstrong NA, Heberling T, Fisher JM, Moore JH. GAMETES: a fast, direct algorithm for generating pure, strict, epistatic models with random architectures. BioData Min. 2012;5:16.

V. Weakening Main Effects and Creating Interactions

Note: Supplementary Files S2 and S4 contain R scripts that we used to generate interactions.

The first step is to select two features in which an interaction is to be created. A proportion of the feature (proportion of instances) are selected to produce a desired strength. The rest of the instances are shuffled to “break” or weaken any main effects.

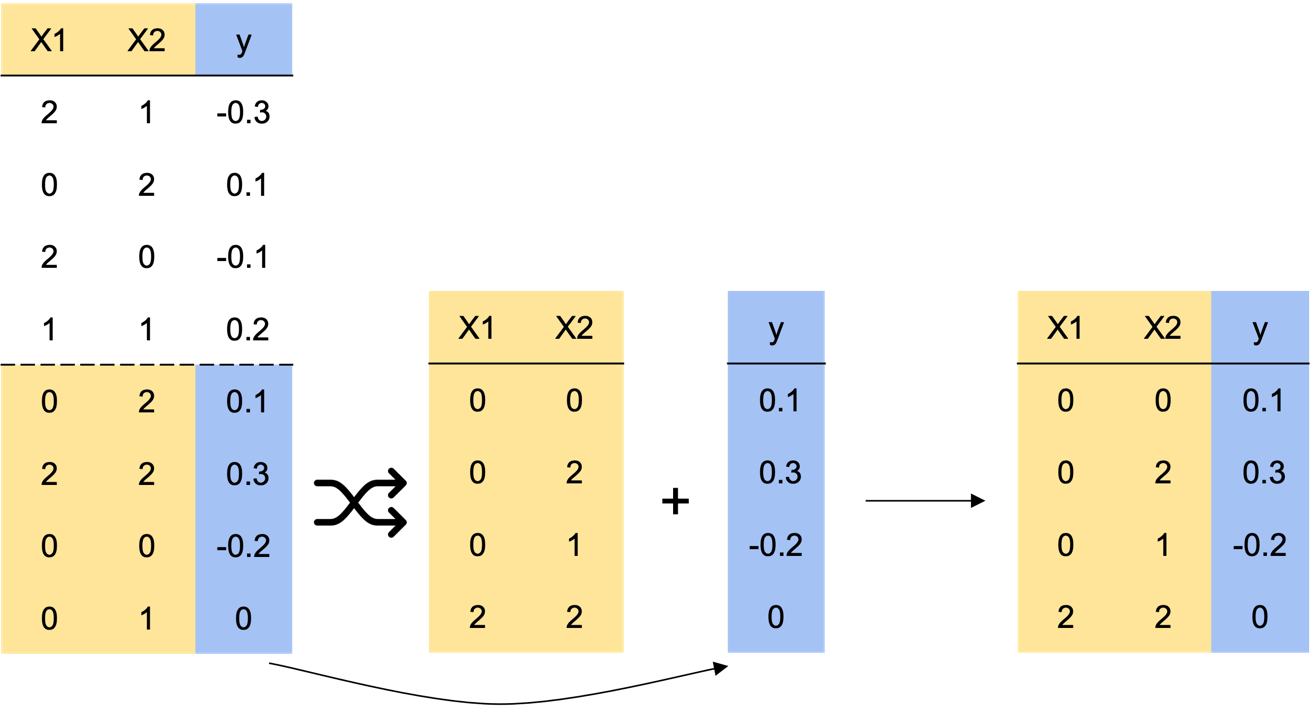

Next, the phenotype instances for the interaction portion are ranked, in this case, in ascending order with the most negative values given the highest rank.

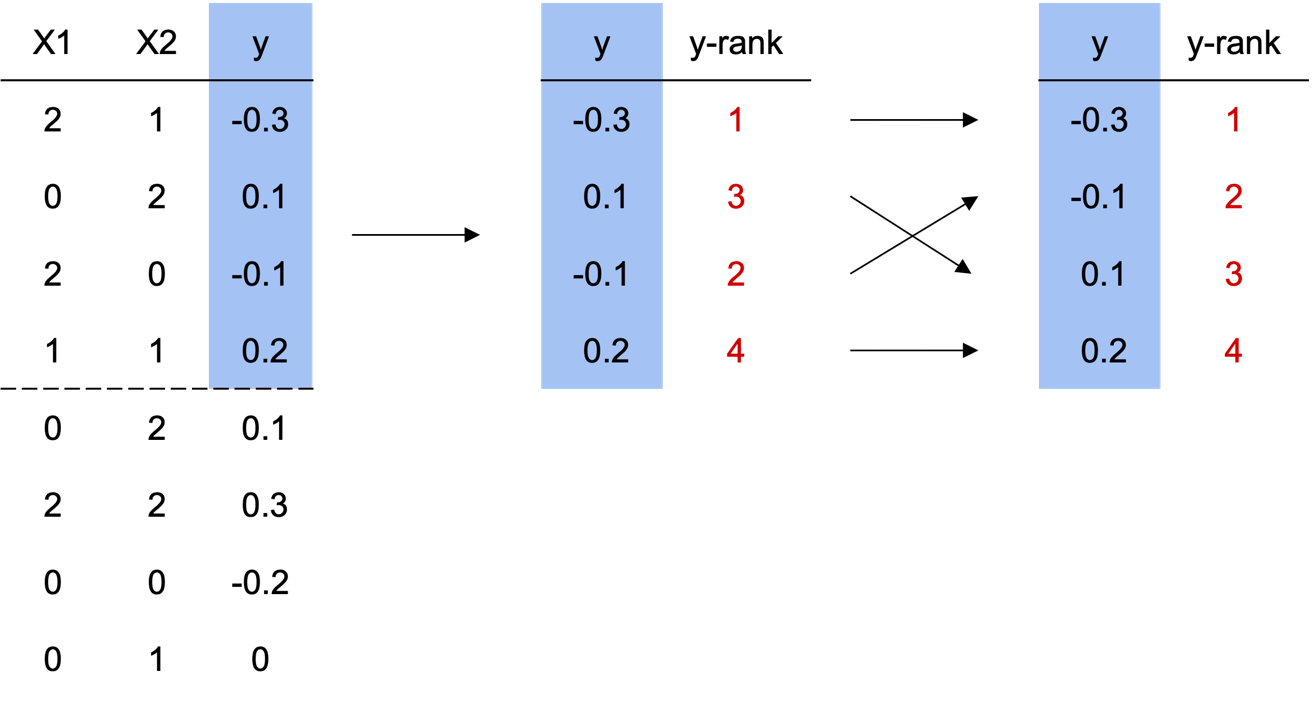

After this, the feature values are shuffled and the XOR penetrance function is created for each pairwise combination. In this example, the combination of 2,2 results in a penetrance function of 0. Then the features, along with their corresponding XOR penetrance, are ranked in ascending order (0’s on top of 1’s).

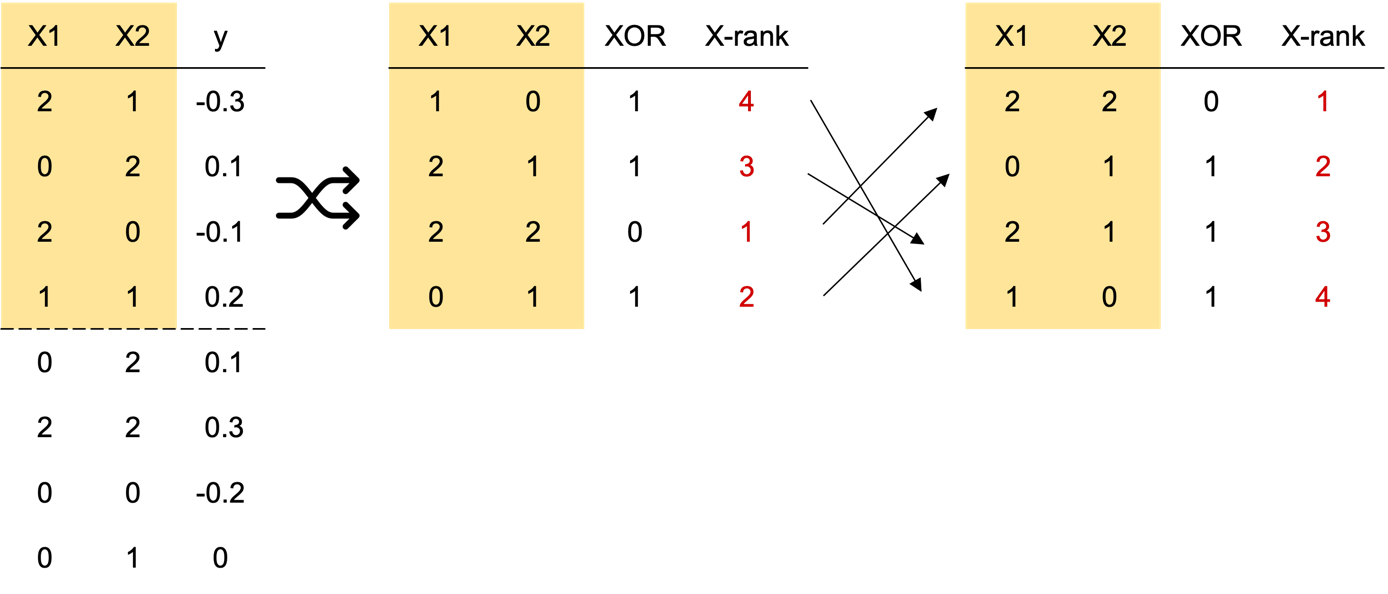

Similarly ranked X’s and y’s are then combined to generate an interaction signal. The XOR feature is removed as it is no longer needed.

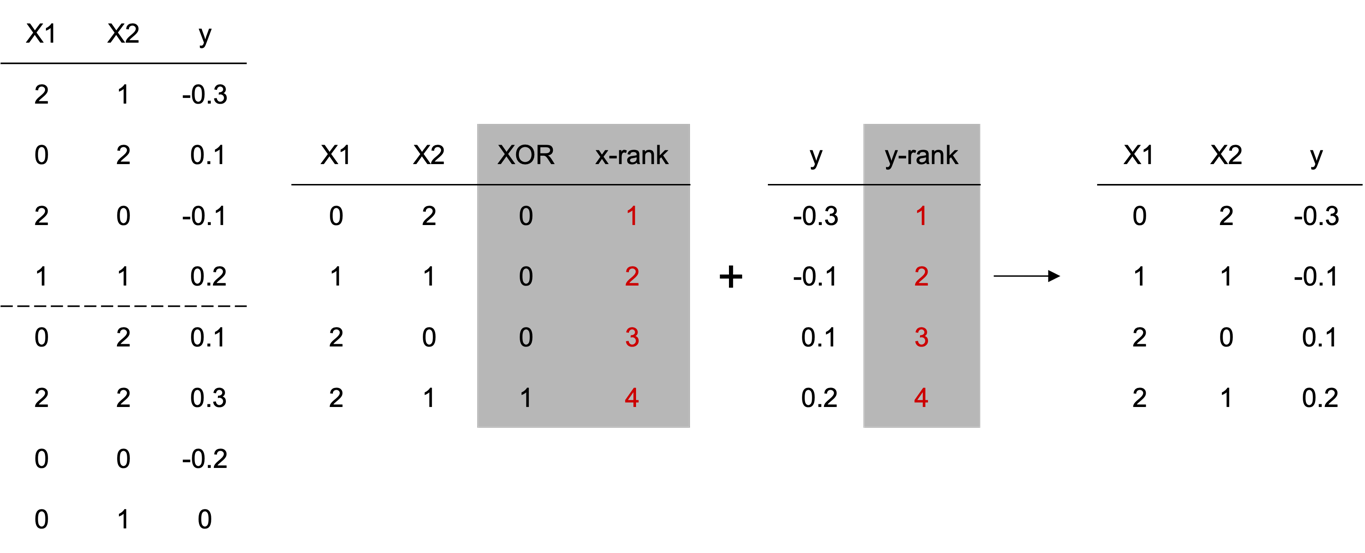

The product are two features that have had their original main effects removed or weakened and an interaction signal generated. The relative strength of the interactions can be altered by selecting less of the instances of the two features to be randomly shuffled/more instances of the two features being maximized for the interaction signal.

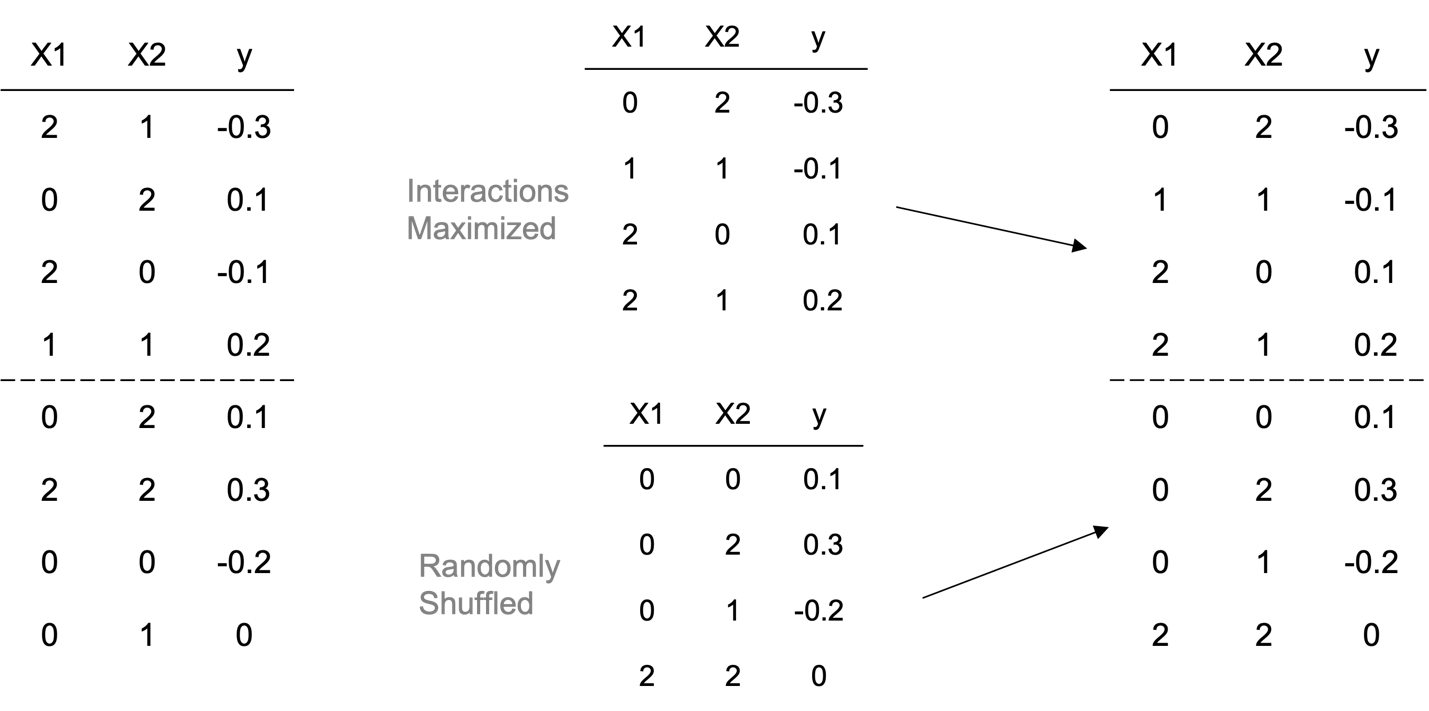

VI. Original QTL p-values (-log_10_P) from GWAS (Chitre et al. 2020, 2022) and linear model containing only putative QTL (performed in R). The last column are the average Shapley values ± S.E.M. obtained for an example run (random seed = 12) in AutoQTL for each locus.

| **SNP (in order of GWAS -log_10_P)** | **-log_10_P (GWAS)** | **-log_10_P (LM with only 18 QTL from R)** | **Avg. Shapley Values ± S.E.M.**  **(AutoQTL)** |
| --- | --- | --- | --- |
| chr1:281788173_G | 21.34521241 | 16.69897 | 0.107 ± 0.0073 |
| chr10:84091208_T | 11.90730391 | 15.69897 | 0.0603 ± 0.016 |
| chr7:129118847_C | 7.875059092 | 7.27876663 | 0.0596 ± 0.0018 |
| chr5:107167969_G | 7.509347766 | 6.27075419 | 0.0353 ± 0.0088 |
| chr19:24321261_T | 7.501719779 | 8.13780897 | 0.0324 ± 0.0081 |
| chr4:178946041_A | 7.443924357 | 9.17966716 | 0.0363 ± 0.0095 |
| chr10:23267180_G | 7.121737365 | 5.18482087 | 0.0262 ± 0.011 |
| chr7:8599340_A | 7.079819781 | 5.1932775 | 0.0285 ± 0.012 |
| chr1:203085725_C | 6.918515871 | 9.99139983 | 0.0367 ± 0.013 |
| chr3:136492861_G | 6.871180858 | 11.8880657 | 0.0466 ± 0.012 |
| chr18:27348077_G | 6.822974566 | 4.56177419 | 0.0406 ± 0.011 |
| chr18:32316331_A | 6.746695925 | 5.83238733 | 0.00699 ± 0.0070 |
| chr5:72916242_T | 6.350971982 | 4.39848322 | 0.0366 ± 0.011 |
| chr2:241577141_C | 6.275761826 | 10.1201589 | 0.0435 ± 0.013 |
| chr9:71715296_A | 6.273175025 | 4.59670786 | 0.00553 ± 0.0055 |
| chr9:15866960_A | 6.113209884 | 6.12453358 | 0.0378 ± 0.0096 |
| chr6:29889998_C | 5.658686185 | 9.98005332 | 0.0610 ± 0.0014 |
| chr8:103608382_G | 5.614770994 | 5.50182734 | 0.0210 ± 0.0079 |

References:

Chitre, A. S. *et al.* Genome-Wide Association Study in 3,173 Outbred Rats Identifies Multiple Loci for Body Weight, Adiposity, and Fasting Glucose. *Obesity* **28**, 1964–1973 (2020).

Chitre, A. S. *et al.* Genome-Wide Association Study in 3,173 Outbred Rats for Body Weight, Adiposity, and Fasting Glucose. *Genes and Addiction: NIDA Center for GWAS in Outbred Rats* <https://cgord.org/dataset/2> (2022).

R Core Team. R: A language and environment for statistical computing. (2022).

VII. The evolution of the average scoring metrics over the course of six generations of GP optimization for the 18 QTL dataset.

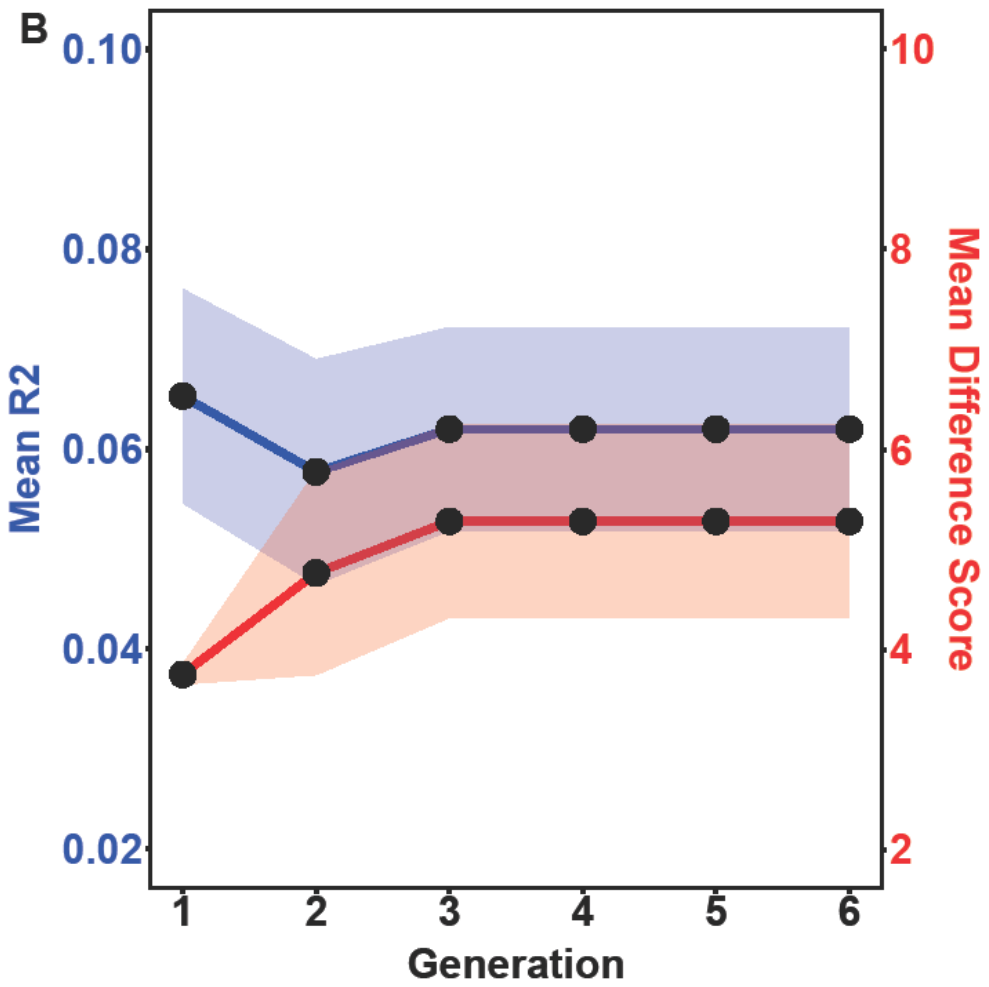

Mean R^2^ (blue line with blue shaded S.E.; left axis) and Mean DS (red line with red shaded S.E.; right axis) over 6 GP generations for an 18 QTL dataset example run (AutoQTL random seed split = 12).

VIII. The evolution of the average scoring metrics over the course of 25 generations of GP optimization for the XOR dataset.

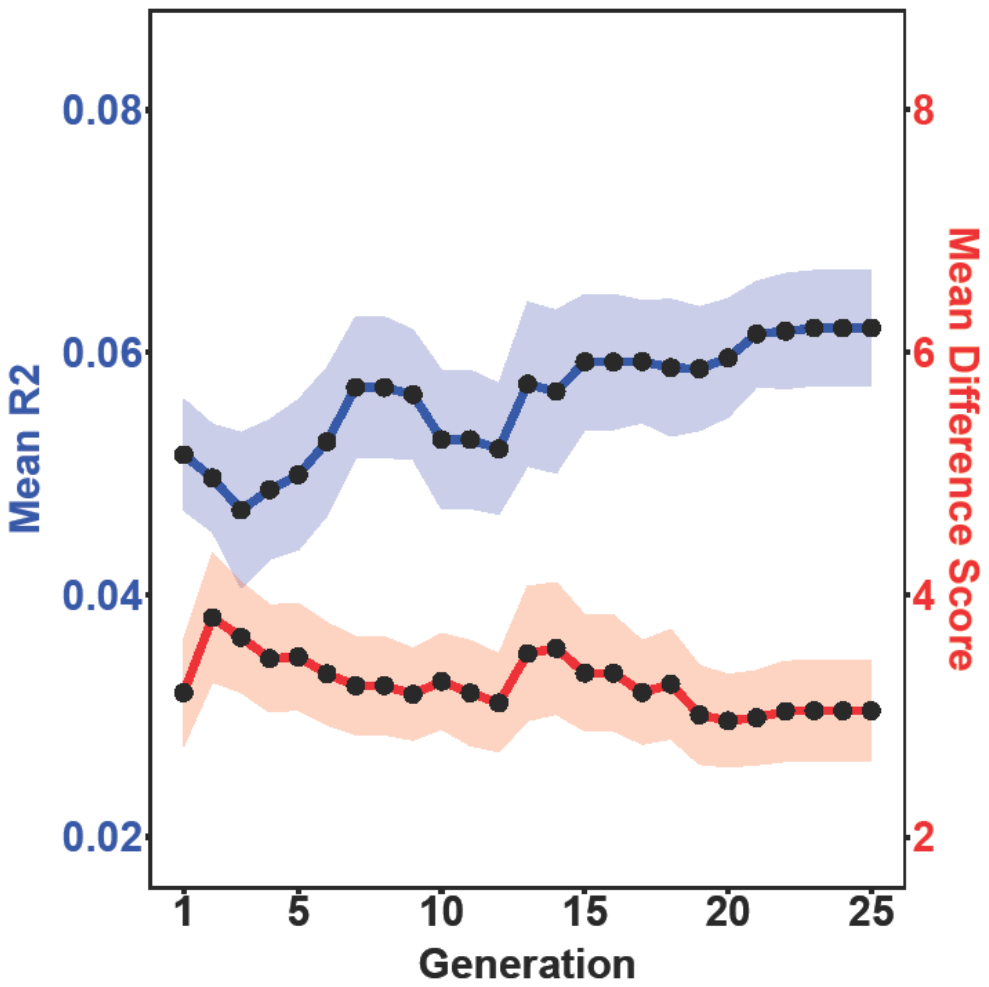

Mean R^2^ (blue line with blue shaded S.E.; left axis) and Mean DS (red line with red shaded S.E.; right axis) over 25 GP generations for an XOR dataset example run (AutoQTL random seed split = 12).

IX. Average SHAP feature importance scores across 29 pipelines from the example AutoQTL run from the XOR dataset for each locus.

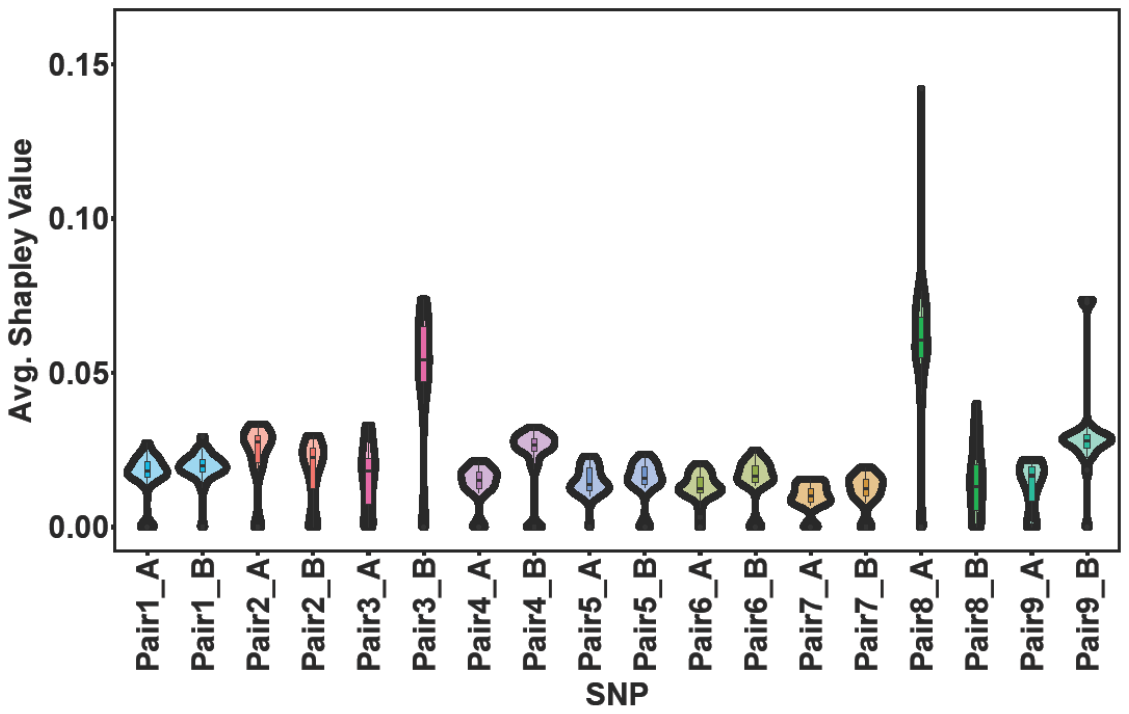

Violin plots and boxplots (inside violin plots) of the same color denote interaction pairs.

X. Multiple linear regression adjusted R^2^ comparison for each of the 18 putative QTL under the six different inheritance models used in AutoQTL. These adjusted R^2^ values were obtained from multiple linear models after each locus was changed to a different inheritance encoding one at a time in R using the code:

Swap[, col:col][Swap[, col:col] == x] <- y

Where “col” is the column currently being re-encoded, “x” is the original encoding for a specific genotype score, and “y” is the new desired encoding of the genotype score. Note: since this procedure re-encodes one genotype state at a time, it is possible to have multiple genotypes have the same score. This is undesirable for 3-level encoders (underdominance and overdominance). Because of this, some genotypes must temporarily be encoded with a placeholder. An example is below for underdominance:

Swap[, 1:1][Swap[, 1:1] == 0] <- 3

Swap[, 1:1][Swap[, 1:1] == 1] <- 0

Swap[, 1:1][Swap[, 1:1] == 2] <- 1

Swap[, 1:1][Swap[, 1:1] == 3] <- 2

After re-encoding is achieved, a multiple linear model is run and the adjusted R^2^ is extracted:

mod <- lm(y ~ ., data = df)

summary(mod)$adj.r.squared

Cells in yellow represent the inheritance model that resulted in the highest model R^2^ for each locus. Cells in light green have a higher model R^2^ compared to additive but are not the highest inheritance model for that locus.

| **SNP (in genomic order)** | **Additive adj. R^2^** | **Dominant adj. R^2^** | **Recessive adj. R^2^** | **Overdominant adj. R^2^** | **Underdominant adj. R^2^** | **Heterosis adj. R^2^** |
| --- | --- | --- | --- | --- | --- | --- |
| chr1:203085725_C | 0.09383 | 0.09362 | 0.09009 | 0.09232 | 0.08871 | 0.09062 |
| chr1:281788173_G | 0.09383 | 0.0837 | 0.09577 | 0.08049 | 0.09134 | 0.08523 |
| chr2:241577141_C | 0.09383 | 0.09261 | 0.09116 | 0.09118 | 0.08841 | 0.08972 |
| chr3:136492861_G | 0.09383 | 0.09265 | 0.08984 | 0.08836 | 0.08652 | 0.08602 |
| chr4:178946041_A | 0.09383 | 0.09324 | 0.09035 | 0.09091 | 0.08812 | 0.08888 |
| chr5:72916242_T | 0.09383 | 0.09315 | 0.09197 | 0.09209 | 0.09042 | 0.09107 |
| chr5:107167969_G | 0.09383 | 0.09261 | 0.09254 | 0.09096 | 0.09062 | 0.09015 |
| chr6:29889998_C | 0.09383 | 0.09267 | 0.09181 | 0.08934 | 0.09002 | 0.08896 |
| chr7:8599340_A | 0.09383 | 0.09254 | 0.09312 | 0.09122 | 0.09260 | 0.09205 |
| chr7:129118847_C | 0.09383 | 0.09262 | 0.09210 | 0.08922 | 0.09052 | 0.08932 |
| chr8:103608382_G | 0.09383 | 0.09394 | 0.09240 | 0.09362 | 0.09233 | 0.09311 |
| chr9:15866960_A | 0.09383 | 0.09199 | 0.09326 | 0.09037 | 0.09125 | 0.09011 |
| chr9:71715296_A | 0.09383 | 0.09392 | 0.09162 | 0.09396 | 0.09401 | 0.09398 |
| chr10:23267180_G | 0.09383 | 0.09266 | 0.09338 | 0.09208 | 0.09306 | 0.09270 |
| chr10:84091208_T | 0.09383 | 0.08953 | 0.09263 | 0.08595 | 0.08910 | 0.08637 |
| chr18:27348077_G | 0.09383 | 0.09240 | 0.09907 | 0.09284 | 0.09669 | 0.09409 |
| chr18:32316331_A | 0.09383 | 0.09379 | 0.09242 | 0.09371 | 0.09330 | 0.09359 |
| chr19:24321261_T | 0.09383 | 0.09366 | 0.09223 | 0.09223 | 0.09112 | 0.09120 |

References:

R Core Team. R: A language and environment for statistical computing. (2022).
